# An evolutionary trade-off between parasite virulence and dispersal at experimental invasion fronts

**DOI:** 10.1101/2020.01.31.928150

**Authors:** Louise Solveig Nørgaard, Giacomo Zilio, Camille Saade, Claire Gougat-Barbera, Matthew D. Hall, Emanuel A. Fronhofer, Oliver Kaltz

## Abstract

Eco-evolutionary processes may play an important role in the spatial spread of infectious disease. Current theory predicts more exploitative parasites to evolve in highly connected populations or at the front of spreading epidemics. However, many parasites rely on host dispersal to reach new populations. This may lead to conflict between local transmission and global spread, possibly counteracting selection for higher virulence. Here, we used the freshwater host *Paramecium caudatum* and its bacterial parasite *Holospora undulata* to investigate parasite evolution under an experimental range expansion scenario with natural host dispersal. We find that parasites evolving at experimental range fronts favoured higher dispersal rates of infected hosts than did parasites evolving in core populations. Front parasites further showed lower levels of virulence (host division and survival) and delayed development of infection, consistent with parameter estimates from an epidemiological model that we fitted on experimental time-series data. This combined evidence suggests an evolutionary trade-off between virulence and host-mediated dispersal, with a concomitant reduction in the investment into horizontal transmission. Our experiment illustrates how parasite evolution can be shaped by divergent selection encountered in different segments of an epidemic wave. Such an interplay between demography and spatial selection has important implications for the understanding and management of emerging diseases, and, more generally, for biological invasions and other non-equilibrium scenarios of spreading populations.

**SIGNIFICANCE STATEMENT:** What drives parasite evolution in spatially expanding epidemics? Many parasites require dispersal of infected hosts to reach new patches, and this may produce specific adaptations enhancing spatial spread. We performed experimental range expansions in an aquatic model system, with natural dispersal of infected hosts. Parasites from experimental range fronts were less virulent and interfered less with host dispersal, but also invested less in horizontal transmission than parasites from the range core. Thus, dispersal adaptation at the front may come at a cost of reduced horizontal transmission, a trade-off rarely considered in theoretical models on parasite virulence evolution. These results have important implications in the context of emerging diseases, and for parasite evolution during biological invasions or other spatial non-equilibrium scenarios.

## INTRODUCTION

In an increasingly connected world, and with changing environments and habitats, we are facing the risk of infectious diseases spreading outside their natural range and over large geographic scales (1–5). This issue is of concern to human health, agriculture and wildlife conservation, and understanding the ecological and evolutionary drivers represents a major challenge to epidemiologists and evolutionary biologists (6–8). Due to their short generation time and large population sizes, parasites have the potential to evolve rapidly, and therefore one important question is whether changes in transmissibility or virulence already occur while an epidemic is progressing (9). Classic theory predicts evolutionary optima for these traits only in large, spatially homogeneous populations at equilibrium (10, 11), but these conditions are unlikely to be met during an epidemic (12, 13). In patchy real-world populations, parasites experience extinction-recolonization dynamics typical of metapopulations, with epidemic spread critically depending on population connectivity and the mobility and dispersal of infected hosts (14–19). Although fundamental for epidemiology, it is still unclear how these spatio-temporal aspects affect concomitant evolutionary processes, and whether they might even lead to specific parasitic adaptations enhancing the spatial spread of the epidemic (see 20–22)

Recent theory has begun to develop a conceptual framework to investigate parasite evolution in spatially explicit, non-equilibrium settings (23). Assuming a classic virulence-transmission trade-off (24, 25) and local feedbacks between epidemiology and selection, several models predict that more virulent parasites will evolve in highly connected “small-world” landscapes (26–28) or at the front of advancing epidemics (20), where host exploitation and transmission is not limited by local depletion of susceptible hosts (“self-shading”). These predictions are consistent with observed changes in the predominance of a highly virulent honeybee virus at the front of progressing epidemics in New Zealand (29), and potentially also with observations for parasites and pathogens of amphibian species (30, 31).

Yet, not all host-parasite systems show this pattern. For instance, the geographic spread of a bacterial pathogen of North American house finches was associated with decreased virulence in the newly invaded areas (32). Likewise, in monarch butterflies, hosts that sustain long or frequent migrations were found to harbour less virulent parasites (33). This may be explained by the way parasite dispersal enters into the equation. Namely, if parasites travel with their infected hosts, exploitation of host resources may reduce dispersal, thereby introducing a novel dispersal-virulence trade-off. Osnas et al. (2015) (22) show that such trade-off can lead to selection favouring more prudent and dispersal-friendly parasites at the moving edge of an epidemic that escape more virulent and transmissible parasites from the core of an epidemic. This latter idea mirrors classic principles from metapopulation theory and metacommunity ecology, based on trade-offs between competitive ability and colonisation/dispersal (20, 34–36). It also relates to recent work on invasive species and range expansions, where dispersal evolution plays a key role in determining the rate of spatial diffusion (37). In this sense, parasites may evolve ‘invasion syndromes’, with characteristic changes in morphology, life history or transmission strategies (30, 31, 38), thereby creating a positive feedback loop between rates of dispersal and rates of spatial spread of infection.

Although the study of naturally expanding parasites remains the ultimate litmus test of the theory, controlled experiments can verify important assumptions and serve as proof of principle (39). For example, we can manipulate demographic conditions in experimental microcosms to mimic the front and core of an expanding epidemic (36) or artificially change levels of population mixing to study epidemiological or evolutionary processes (39). Indeed, studies of the latter type found that experimentally shifting populations from local to global “dispersal” favoured more virulent parasites (40–42), as predicted by theory (23, 43). Yin (1993) (44) further showed that phage diffusion on bacterial lawns is associated with the appearance of faster replicating mutants in the periphery. However, to our knowledge, there are no studies addressing experimental evolution of parasites from an explicit metapopulation perspective, under natural dispersal of a host and its parasite.

For (micro-)organisms with directed movement, experimental landscapes can be created to study metapopulation processes or range expansion dynamics with natural dispersal (45–47). Here, we employed such an approach to investigate the experimental evolution of spatially spreading parasites, where all parasite dispersal is host-mediated. Using two-patch dispersal arenas for the ciliate *Paramecium caudatum* infected with the bacterial parasite *Holospora undulata*, we mimicked a range expansion scenario, with a front population of hosts (and parasites) dispersing into a new microcosm during each selection event, and a core population constantly remaining in place (and losing emigrants; see Fig. 1). After 55 episodes of dispersal selection, we then assayed evolved front and core parasites under common garden conditions on naive hosts. Multiple traits were measured, namely the parasites’ effect on host dispersal, investment in horizontal transmission and their impact on host replication and survival. We further obtained additional independent estimates of parasite traits by fitting a simple epidemiological model to time series data (population density, infection prevalence) from the experimental assay.

**Fig. 1.**
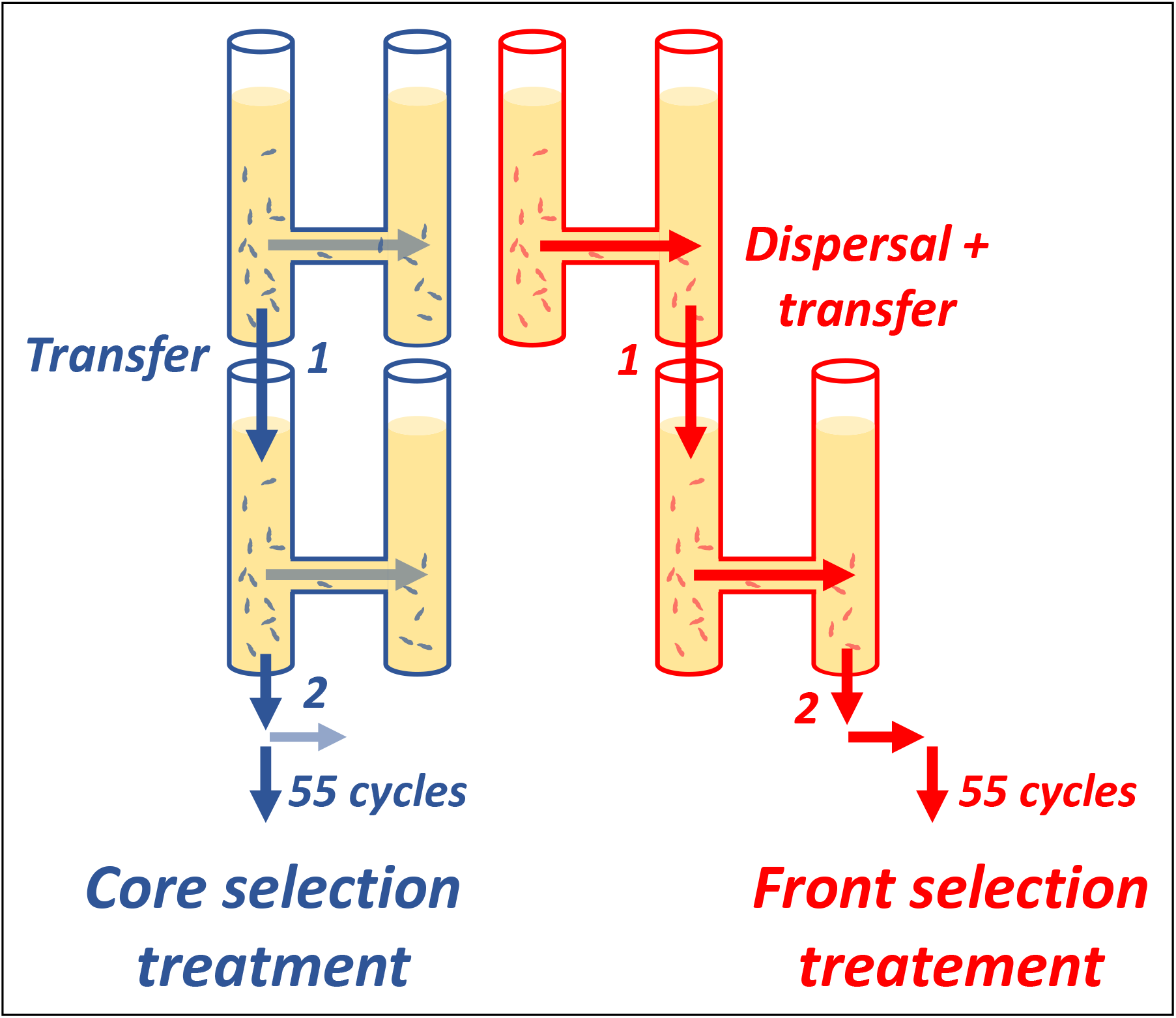
Experimental design of the long-term selection experiment, using 2-patch dispersal systems. Infected populations were placed in the ‘core’ tube and allowed to disperse to the other ‘front’ tube during 3 h (horizontal arrows). In the front selection treatment (red), only the dispersing fraction of the population was maintained, whereas in the core selection treatment (blue) only the non-dispersing fraction was maintained. After adjustment of initial densities, the selected fractions were then transferred to a new tube (vertical arrows) and grown for 1 week, during which time demographic and epidemiological processes acted freely. A total of 55 dispersal/growth cycles were performed, for 5 core and 5 front selection lines.

Because parasite persistence in the front populations depended entirely on host dispersal, we predicted that front parasites would evolve minimal impact on host dispersal, or even increase dispersal of infected hosts (48). Such dispersal adaptations could involve a decrease in parasite virulence (22) and generate an evolutionary trade-off with investment in horizontal transmission, not expected to occur in the core populations. Our results were broadly consistent with these predictions, and we conclude that differential dispersal selection pressures arising at the core and front of a range expansion can lead to marked divergence of parasite life-history traits and the emergence of a ‘parasite dispersal syndrome’.

## RESULTS

Evolved parasites from the five front and five core selection lines were extracted and the inocula used to infect naïve hosts (three genotypes). Several traits were measured for these newly infected assay cultures (for timing of assays see Table S1 and statistical analyses Tables S2, S3).

### Infected host dispersal

On average, hosts infected with front parasites dispersed twice as much (mean dispersal rate: 0.24 ± 0.05 SE × 3h^−1^) as those infected with the core parasites (0.12 ± 0.02 SE; Fig. 2A). This effect of selection treatment was significant (χ^12^ = 4.9, p = 0.027; Table S2). The distribution of the differences between model predictions for front and core treatments (small panel, Fig. 2A) also shows that higher front-parasite dispersal is the most frequent predicted outcome (>98%; mean front-core difference: 0.12, 95% CI [0.08; 0.35]). This general trend was consistent on all three host genotypes tested (Fig. S4; Table S3). Fig. S4 further shows that levels of front-parasite dispersal were similar to reference data for uninfected hosts, whereas core parasites generally reduced dispersal.

**Fig. 2.**
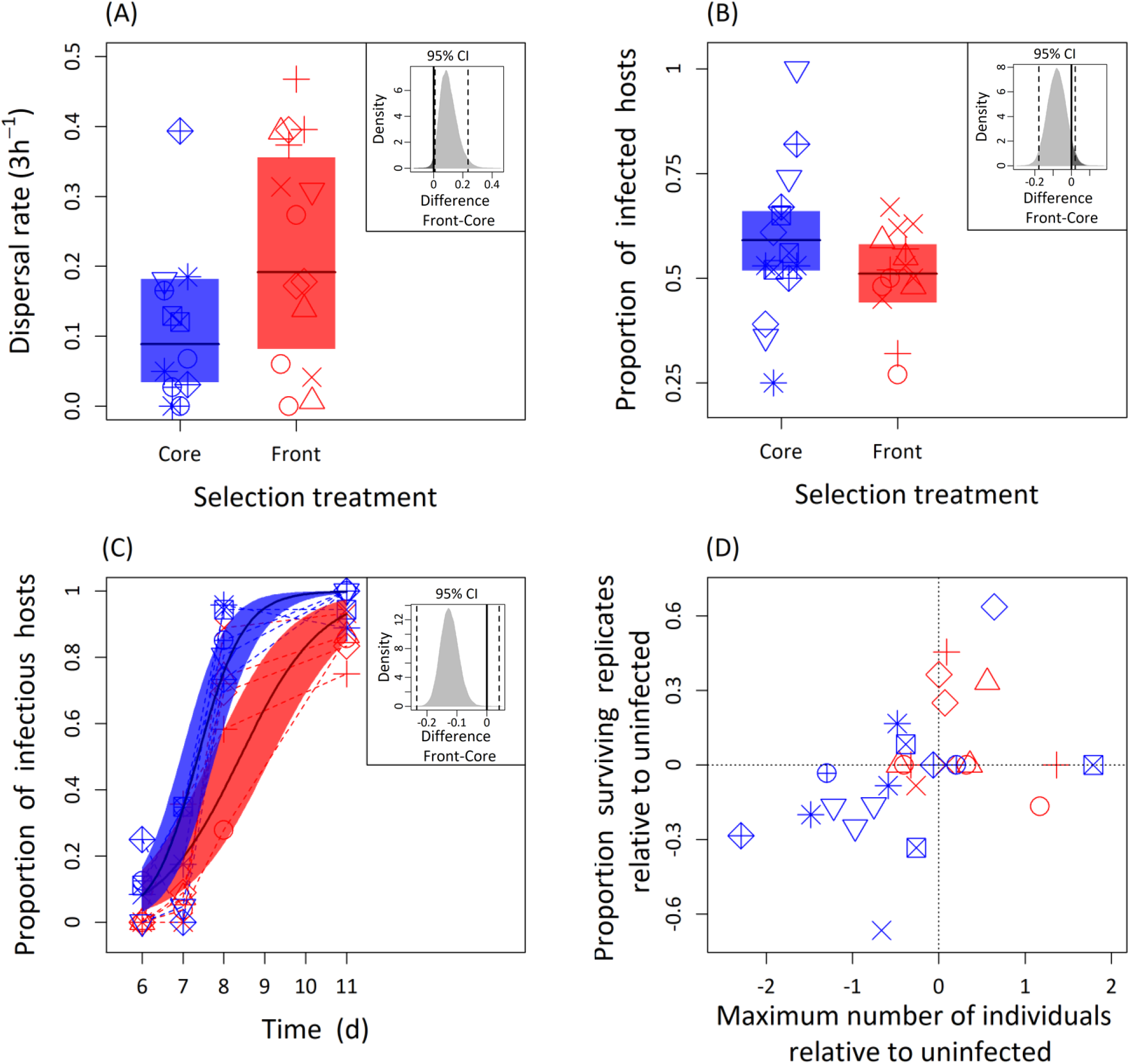
Dispersal and infection life-cycle traits of evolved parasites from core selection (blue) and front selection (red) treatments, measured on naïve *Paramecium*. **(A) Dispersal rate**. Proportion of dispersing infected hosts observed in infected assay cultures placed for 3 h in a dispersal system. **(B) Infectivity.** Proportion of infected hosts in assay cultures on day 4 post-inoculation (p.i.). **(C) Infectiousness**. Proportion of infectious hosts in infected assay cultures between day 6 and 11 p.i.. Infectious hosts are individuals that produce infective spores of the parasite. **(D) Virulence.** Association between infected host division and survival, expressed relative to uninfected hosts (infected minus uninfected). Negative values indicate negative effects of infection on the host trait. Panels (A)-(C) show means and 95 % confidence intervals of the model predictions. Small insert panels show predictive distributions (and 95% CI) of the difference between front and core treatments. Symbols represent observed means for different combinations of parasite selection line and assay host genotype. Different symbols refer to different parasite selection lines (n = 10).

Using video analysis, we investigated variation in two parameters of *Paramecium* swimming behaviour: swimming speed and trajectory variation (tortuosity). We found no evidence that infection with core or front parasites had significant effects on these two parameters (p > 0.25; Table S2, neither when tested on host genotypes individually S3; Fig S5, S6). Moreover, mean levels of swimming speed or tortuosity were not significantly correlated with infected dispersal rates (r ≤ 0.15, n = 29, p > 0.4), indicating that dispersal was not directly affected by these aspects of swimming behaviour (see also path analysis below).

### Parasite life-history traits

#### Infectivity

Measurements of infection prevalence early after inoculation (day 4) inform on parasite horizontal transmission potential (infectivity). Core parasites had slightly higher infection success (selection line average proportion of infected hosts: 0.59 ± 0.05 SE) than front parasites (0.51 ± 0.03 SE; Fig. 2B). Although not formally significant (effect of selection treatment: χ^12^ = 2.43, p = 0.118; Table S2; predictive difference distribution front-core infectivity: mean = −0.08, CI [0.02; −0.18]; Fig 2B; see also Fig. S7 for genotype specific responses), this trend was consistent with higher estimates of the transmission parameter for core parasites in an epidemiological model fitted to our experimental data (see below; Fig. 4).

#### Investment in horizontal transmission (infectiousness)

It takes several days until infected hosts produce horizontal transmission stages and become infectious. In our assay, the first infectious hosts appeared on day 6 p.i., and their frequency then increased over the following week, reaching up to 100% (Fig. 2C). Over this period, populations infected with front parasites produced a lower proportion of infectious hosts (mean: 0.41 ± 0.03 SE) than did populations infected with core parasites (0.53 ± 0.03 SE; effect of selection treatment: χ^12^ = 13.2, p <0.001; Table S2; predictive difference distribution front-core infectiousness: mean = −0.10; CI [0.04; −0.23]; Fig. 2C). There was also a difference in timing: on average, core parasites produced the first infectious hosts c. 1 day earlier than did front parasites (day 6 vs day 7) and subsequently showed a faster increase in the proportion of infectious hosts (treatment × time interaction: χ^12^ = 13.54, p < 0.001,Table S2; Fig. 2C). These differences in total investment and/or timing broadly hold on all three host genotypes tested (Table S3; Fig. S8).

#### Virulence

We isolated single infected individuals from the core and front infected assay cultures and measured the impact of infection on host division and survival over a 20-day period. Exposed, but uninfected, controls were isolated from the same assay cultures and run in parallel.

##### Host division

By day 10, 87% of the infected singletons had accomplished at least one division (266 / 305 replicates; mean maximum cell number observed over this period: 8.5 ± 0.6 SE). Analysis of maximum cell density revealed a significant selection treatment x infection status interaction (χ^12^ = 16.9, p > 0.001; Table S2). Namely, hosts infected with front parasites reached nearly twice as high maximum densities (10.9 ± 1.3 SE) than those infected with core parasites (5.9 ± 0.9 SE; contrast front vs core: t_601_ = 4.7, p < 0.0001; predictive difference distribution front-core: mean = 4.6, CI [1.4; 9.6]; Fig. 3A).

**Fig. 3.**
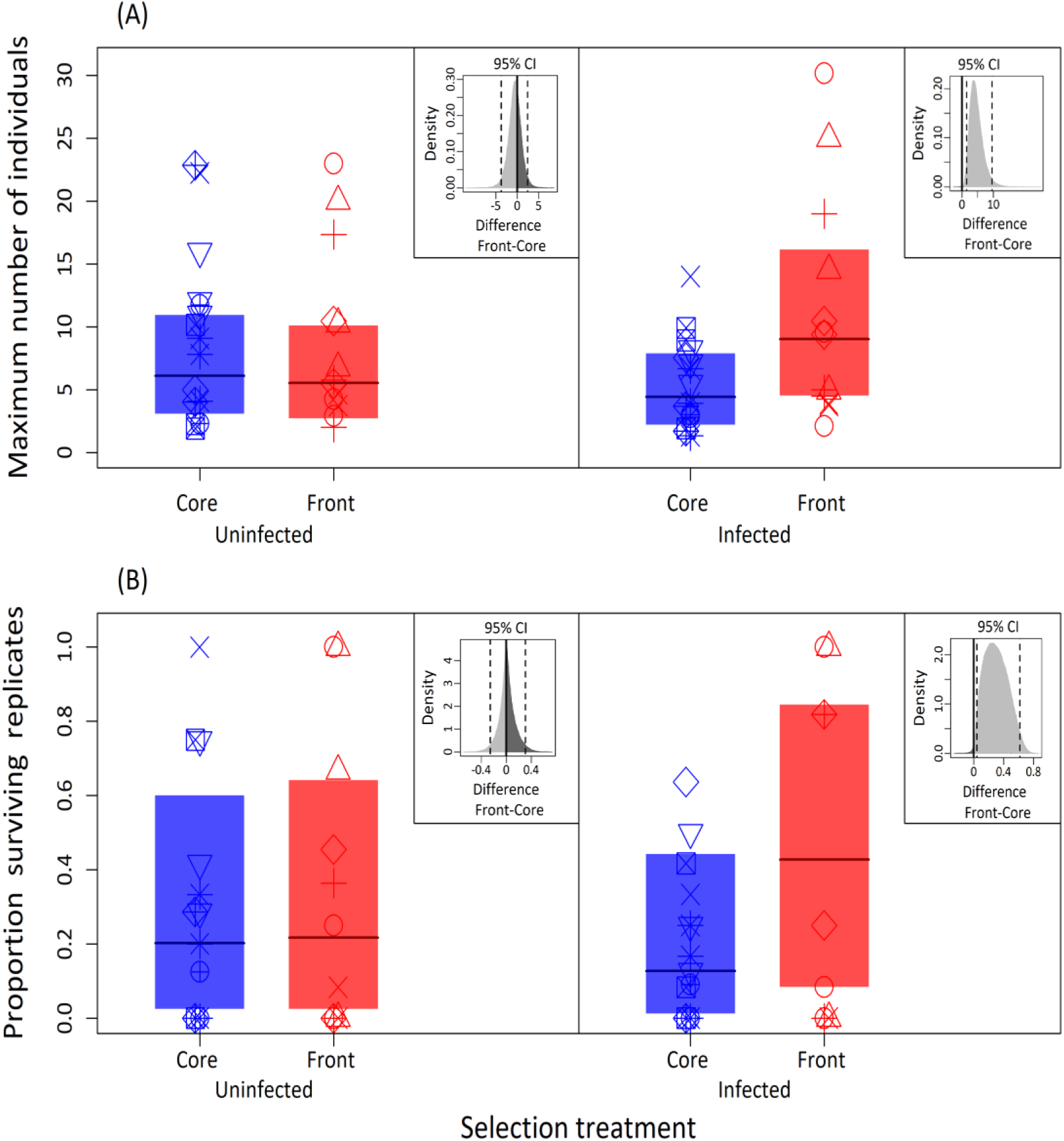
Estimates of virulence of evolved parasites from core selection (blue) and front selection (red) treatments, measured on naïve *Paramecium*. **(A) Host division**. Maximum cell density of infected and uninfected *Paramecium*, as determined in a singleton assay. **(B) Host survival.** Proportion of surviving infected and uninfected replicates on day 20 in the singleton assay. All panels show means and 95 % confidence (CI) intervals of the model predictions. Small insert panels show predictive distributions (and 95% CI) of the difference between front and core treatments. Symbols represent observed means for different combinations of parasite selection line and assay host genotype. Different symbols refer to different parasite selection lines (n = 10).

##### Host survival

As for host division, there was a significant selection treatment × infection status interaction for host survival (χ^12^ = 7.4, p = 0.006; Table S2). By day 20, infections with front parasites had experienced a 50% lower mortality (mean proportion of infected replicates extinct: 0.19 ± 0.05 SE) than infections with core parasites (0.37 ± 0.1 SE; contrast front vs core: t_601_ = 3.75, p > 0.001; predictive difference distribution front-core: mean = 0.30, CI [0.04;0.62]; Fig. 3B). Moreover, effects on host division and on host survival were positively correlated: parasites which allowed more host division also allowed higher host survival (means per parasite selection line: r = 0.84 ± 0.19, n = 10, p = 0.003). Thus, core parasites generally had negative effects, whereas front parasites only had little, or even positive, impact on their hosts reproduction and survival (Fig. 3A and 3B), and these opposing trends were consistent across the three host genotypes tested (Table S2; Fig. S9 & S10).

### Path analysis

Using path analysis, we explored the direct and indirect contributions of different traits to the observed variation in infected host dispersal (Fig. 4A). Host division (= maximum cell density) was the only trait with a significant direct effect on host dispersal (F_1,20_ = 6.16, p = 0.022; Fig. 3B); thus, lower virulence was associated with higher dispersal rates of infected hosts. Horizontal transmission investment (= cumulative infectiousness) had a moderate indirect effect on dispersal via its significant negative relationship with virulence (F_1,23_ = 4.47, p = 0.0456; Fig. 3C). Swimming behaviour (speed, tortuosity) had no significant direct effect on dispersal, and were themselves only very marginally affected by virulence or infectiousness (Fig. 4A).

**Fig. 4.**
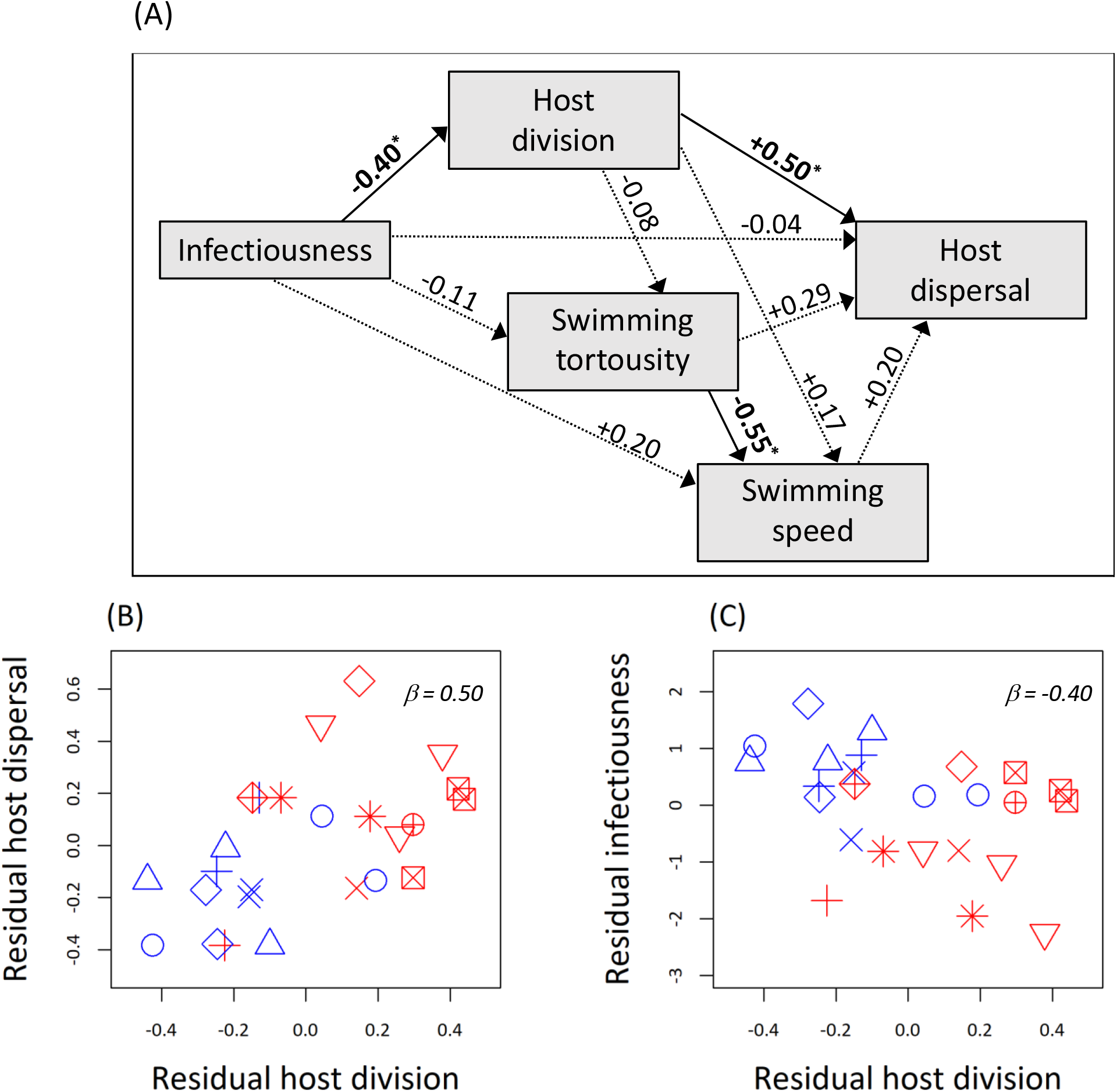
Relationships between parasite traits. **(a)** Path analysis testing direct and indirect effects of 4 parasite traits on infected host dispersal: (i) Infectiousness (cumulative proportion of host producing infective spores; area under the curve: day 6 - 11 p.i.); (ii) Host division (maximum infected cell density); (iii) Swimming tortuosity (≈ trajectory changes) and (iv) net swimming speed of infected singletons. Analysis based on trait means for different combinations of parasite selection line and host assay genotype (n = 25) and performed on residuals, after correcting for overall effects of host assay genotype. Standardised beta regression coefficients (β) are shown above arrows (*p < 0.05); **(b)** Relationship between residual host division and dispersal; **(c)** Relationship between residual horizontal transmission investment and host division.

### Epidemiological model fits

By fitting an epidemiological model to the population-level data from the assay replicate cultures (infection prevalence and population density), we obtained independent estimates of parasite parameters. In the model, we integrated the basic features of the infection life cycle, assuming simple population growth and regulation (Beverton-Holt type model, 49) and parameterising virulence as the reduction in host fecundity.

The model captured the main trends in the demographic and epidemiological dynamics observed in the cultures. This is illustrated in Fig. 5A, showing the model fits for the densities of infected and uninfected hosts for the 63D host genotype (for the other two host genotypes, see SI 4, Fig. S11). Parameter estimates confirm the main trends found in our experimental assays. Namely, the model fits show that front parasites have lower virulence, lower transmission rate and longer latency time than core parasites (Fig. 5B-D), a pattern largely consistent for the three host genotypes tested (Fig. S11).

**Fig. 5.**
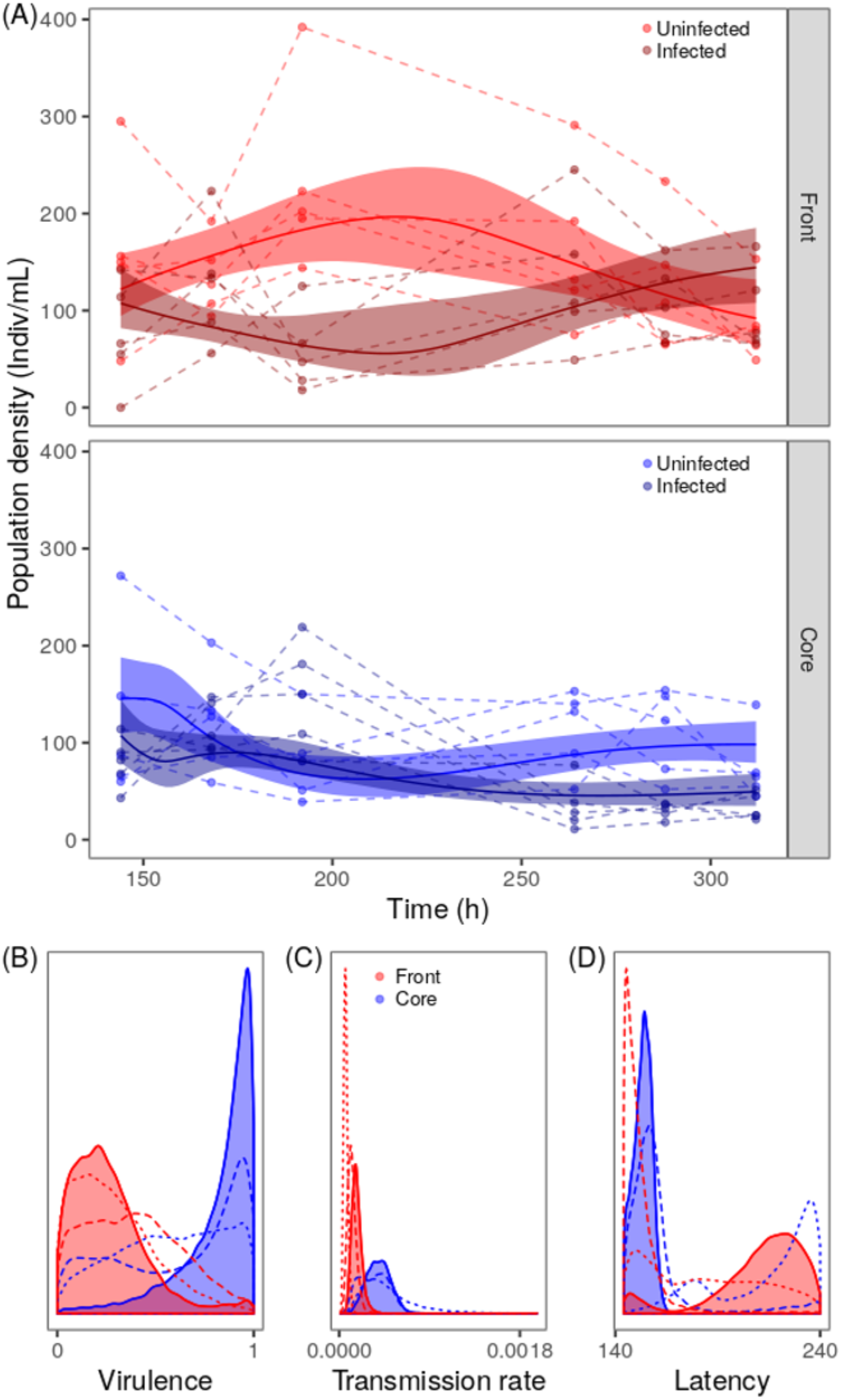
Fit of the epidemiological model. **(a)** Fit of the epidemiological model (equations 6–8) to infected and uninfected host density time-series data, obtained for assay cultures infected with core (blue) and front (red) parasites. Curve fits shown for host genotype 63D (for other genotypes, see Fig. S11). Dashed lines represent observed densities for different replicate assay cultures, solid lines and shaded areas represent posterior model predictions (mean and 95% CI). **(b)-(d)** Posterior distributions for virulence (= reduction in host division rate), horizontal transmission rate and latency, respectively. Solid lines and shaded areas show posterior distributions for host genotype 63D, dashed lines for host genotype C173, and the dotted lines for genotype C023.

## DISCUSSION

In times of global epidemics (1, 4, 32) it is important to know how parasites evolve while spreading through a landscape or entire continents. Recent theory suggests that spatial ‘viscosity’ and connectedness generate eco-evolutionary feedbacks, with important consequences for parasite virulence evolution and the speed of epidemics (23). However, so far little attention has been given to the fact that many parasites travel together with their dispersing hosts, which may considerably affect evolutionary predictions (28, 50, 51).

To address this issue, we performed a simplified range expansion experiment, with natural dispersal of infected hosts. Our ‘range front’ and ‘range core’ treatments imposed differential selection on host dispersal (see 52) and resulted in divergent parasite phenotypes: front parasites allowed for higher dispersal of their infected hosts, were less virulent and showed reduced investment in horizontal transmission, compared to the parasites from the core selection treatment. These patterns were largely robust between the three naive host genotypes tested, and additionally confirmed by results from an epidemiological model that we fitted to time-series data obtained from our assay cultures.

### Evidence for a virulence - dispersal trade-off

Our experimental result of multi-trait changes joins empirical observations of “invasion syndromes” in naturally spreading diseases, such as avian malaria in Europe (53) or lungworms of invasive cane toads in Australia (30). Lungworms at the invasion front, for example, exhibit distinct life-history traits (reduced age at maturity, larger infective and free-living larvae), possibly representing adaptations to invasion history (30). We replayed such an invasion history, by mimicking the spatial progression of an isolated population in our range front treatment, which was expected to favour parasites that succeed in dispersing together with their infected hosts. This explains why our front parasites were found to facilitate higher host dispersal. Importantly, higher host dispersal was associated with higher host replication and survival, indicating a dispersal - virulence trade-off (Fig. 2 and 3). Reduced virulence, in turn, was associated with reduced horizontal transmission potential (Fig. 2C), consistent with previous findings in this system (54–56) and reflects a virulence-transmission trade-off for this parasite. Thus, we conclude that the evolution of higher parasite dispersal in front parasites came at the cost of reduced horizontal transmission, a trade-off resulting from a reduction in virulence.

The idea that parasite exploitation strategies can be shaped by the interplay between local transmission and global dispersal was formalised in a theoretical model by Osnas et al. (2015) (22). They showed that implementing a trade-off between virulence and the capacity of infected hosts to disperse, favours less virulent strains at the front of an epidemic, escaping the more competitive (and more virulent) strains through faster dispersal (22). Such a selection scenario may explain observed geographic patterns of virulence for a bacterial pathogen of North American house finches (32), and it is qualitatively consistent with our results.

### Trait relationships: Proximate causes of infected dispersal rate

Just like parasites can alter their hosts behaviour to increase transmission (57), they may also evolve to manipulate host dispersal (50, 58). However, we find little evidence for manipulation to increase the dispersal. Consistent with previous observations of negative effects of infection in this (59) and other systems (36, 60–62), core parasites reduced host dispersal, whereas infection with front parasites produced levels of dispersal comparable to uninfected *Paramecium*. Path analysis indicates that virulence is the main direct predictor of host dispersal in our assays. Investment in horizontal transmission had an indirect effect via decreased virulence. Although intuitively straightforward through a weakening of infected hosts, the mechanistic link between virulence and dispersal remains unclear. We found no effect of infection on swimming behaviour, nor was there a direct link between swimming behaviour and dispersal, which is frequently observed in other protists (63). Possibly, infection influences other dispersal-relevant traits, such as the vertical distribution in the water column (64, 65), determining the proximity of individuals to the opening that leads to the other tube in the dispersal arena (see Fig. 1).

### Contrasting scenarios: Dispersal to new sites vs access to new hosts

While our results suggest more prudent parasites might be spreading at invasion fronts, other theoretical models and experiments reach opposite conclusions. Griette et al. (2015) (21), for example, predict highest levels of virulence at the front of an epidemic wave, where transmission is not limited by the availability of susceptible hosts, thereby favouring the most ‘rapacious’ variants. Following this line of argument, experiments with viruses and bacteriophages have studied virulence evolution by artificially manipulating dispersal or population connectivity (40–42). Kerr et al. (2006) pipetted bacteria and phage either to adjacent wells or to more distant wells on a multi-well plate, in analogy to our ‘core’ and ‘front’ treatments (41). Contrary to our results, their latter treatment of unrestricted dispersal resulted in the evolution of more virulent phages, confirming the prediction that erosion of spatial structure in ‘small worlds’ favours more transmissible and more virulent parasites (27, 28, 66).

One reason for these contrasting results is that in Kerr et al. (2006) (41), dispersal was artificial and cost-free, eliminating a possible virulence-dispersal trade-off. Secondly, our experiment considered a different spatial scenario where infected hosts disperse into empty space, more characteristic of a biological invasion. This means that higher dispersal was not rewarded with more access to susceptible hosts, as assumed in the above models (20, 66). Future experiments can test whether we still find reduced virulence in the front selection treatment, if infected hosts disperse into patches already occupied by uninfected hosts.

Taken together, these examples illustrate the different ways in which spatial spread and dispersal of parasites can be approached both conceptually and experimentally, with very different evolutionary outcomes. We argue that particular attention should be given to *how* parasites disperse through a landscape, namely because dispersal itself may be the target of selection (36, 51).

### More vertical transmission at invasion fronts?

We used the replication of infected hosts as a measure of virulence. However, for this parasite, host replication is also equivalent to vertical transmission, because reproductive stages are passed on to daughter cells during mitosis. In this sense, parasites in our front selection treatment underwent a shift from horizontal transmission towards higher levels of vertical transmission. This is due to an underlying developmental trade-off, where reduced conversion of reproductive into infective stages decreases the negative effects on host replication (56, 67), but simultaneously reduces horizontal transmission capacity.

Magalon et al. (2010) observed a similar increase in the efficacy of vertical transmission of this parasite in frequently disturbed populations (55). In fact, their study can be re-interpreted as a range expansion experiment, with the disturbance treatment mimicking the frequent recolonization events occurring at the front (68) and the less disturbed control treatment reflecting more stable conditions in the core (see Fig. 1 in 55). The explanation for the evolutionary shift towards vertical transmission is that the demographic oscillations at the invasion front, with frequent periods of low host density and high host fecundity, increase the contribution of vertical transmission to the total transmission success (69). Because vertical transmission is a means of ‘reproductive insurance', we would generally expect it to evolve in association with parasite dispersal syndromes in expanding populations or in highly disturbed habitats (70). We note, however, that in our present experiment both core and front populations went through density bottlenecks, making the dispersal constraint the main selective driver.

### Conclusions

Our results show that different segments of an epidemic wave may be under divergent selection pressures. Namely, we find evidence that dispersal selection at an experimental invasion front leads to reduced virulence. This contrasts with observations in certain natural epidemics (29, 30), while confirming others (32, 33). This calls for more detailed investigations of the role of dispersal for epidemic spread and its implications for the evolution of parasite life-history traits. Our relatively simple statistical modelling exercise suggests that time series data from natural populations represent a useful resource for such a challenge. Establishing a better understanding of the interaction between demography and rapid evolutionary change in spreading populations is crucial for the management of emerging infectious diseases and disease outbreak in the wild, biological invasions and other non-equilibrium scenarios.

## MATERIALS AND METHODS

### Study system

*Paramecium caudatum* is a filter-feeding freshwater protozoan ciliate from still water bodies in the Northern Hemisphere (71). It has a germline micronucleus and a somatic macronucleus. Our cultures are maintained asexually in a lettuce medium with the food bacterium *Serratia marcescen*s at 23°C, allowing 1-2 population doublings per day (72). The gram-negative alpha-proteobacterium *Holospora undulata* infects the micronucleus of *P. caudatum*, and can be transmitted both horizontally (by s-shaped infective spores, 15 μm) upon host death or during cell division, and vertically, when reproductive bacterial forms (5 μm) segregate into daughter nuclei of a mitotically dividing host (73). After ingestion by feeding *Paramecium,* infective spores invade the micronucleus, where they differentiate into the multiplying reproductive forms. After one week, reproductive forms begin to differentiate into infective spores (64, 72). Infection with *H. undulata* reduces host cell division and survival (56) and also host dispersal (59).

### Long-term experiment

Similar to Fronhofer and Altermatt (2015), we imposed dispersal selection in 2-patch microcosm arenas (Fig. 1, see also SI 1), built from two 14-mL plastic tubes, interconnected by 5-cm silicon tubing, which can be blocked using a clamp (see Fig. S1). We define dispersal as the active swimming of *Paramecium* from one microcosm to the other via the connective tuber (i.e., the dispersal corridor).

The experiment was seeded from an uninfected host line (“63D”, haplotype b05) from our laboratory that had been under “core selection” (see below) for three years and shows characteristically low dispersal propensity (O.K., unpublished data). This 63D mass culture was infected with an inoculum of *H. undulata* prepared from a mix of various infected stock cultures (for details, see SI 2). All parasites in this mix originate from a single isolate of *H. undulata* established in the lab in 2001.

In the front selection treatment, we placed *Paramecium* in one tube (“core patch”) and opened the connection for three hours, allowing them to swim into the other tube (“front patch”). *Paramecium* from the front patch were recovered and cultured in bacterised medium, allowing for natural host population growth and parasite transmission. After one week, we imposed another dispersal episode, again recovering only the *Paramecium* from the front patch, and so on. The core selection treatment followed the same alternation of dispersal and growth periods, except that only *Paramecium* from the core patch were recovered and propagated (Fig. 1, and SI 1). We established five infected ‘core selection’ lines and five infected ‘front selection’ lines that were maintained for a total of 55 cycles of dispersal. To minimise potential effects of host (co)evolution, we extracted infectious forms from each selection line after cycle 30, inoculated a new batch of naïve 63D hosts and continued the experiment for another 25 cycles. For details of the experimental protocols, see SI 1.

### Parasite assays

At the end of the selection experiment, we extracted parasites from core and front selection lines to inoculate new, naïve hosts. We then assayed parasite effects on host dispersal, infection life-history and virulence. To obtain a general picture of trait expression, we tested the evolved parasites on naïve 63D hosts (same genotype as used in long-term experiment), as well as on two other genotypes, C023 and C173 (provided by S. Krenek, TU Dresden, Germany). Companion assays of evolutionary adaptations arising in the host are reported elsewhere (52).

All assays were performed on a cohort of infected replicate cultures, over the course of three weeks under common-garden conditions (Table S1). To initiate the cultures, we placed ≈ 5 × 10^3^ cells of a given naïve host genotype in 1.4 mL of bacterised medium in a 15-mL tube, to which we added the freshly prepared inoculum of a given evolved parasite line (≈ 1.5 × 10^6^ infectious spores, on average). On day four post-inoculation (p.i.), when infections had established, we split the cultures into three technical replicates and expanded the volume to 30 mL, by adding bacterised medium. A total of 90 replicate cultures were set up (2 selection treatments × 5 parasite selection lines × 3 host genotypes 3 technical replicates).

### Dispersal of infected hosts

From day 14 to 19 p.i., we assayed dispersal rates of hosts infected with core and front parasites, using linear 3-patch arenas where the *Paramecium* disperse from the middle tube to the two outer tubes (Fig. S2). Arenas were filled with ~2800 individuals in the middle tube and after 3 h of dispersal, we subsampled the middle tube (0.5 mL) and the pooled two outer tubes (3 mL) to estimate the number of non-dispersers and dispersers under a dissecting microscope. Furthermore, from ≈20 arbitrarily picked individuals stained with 1% lacto-aceto orcein (LAO fixation; Fokin and Görtz, 2009) we determined the proportion of infected dispersers and non-dispersers (phase-contrast, 1000x magnification), from which we then calculated the dispersal rate of infected hosts for each replicate culture (number of dispersed infected hosts / total number of infected hosts per 3 h). Each of 88 available replicate cultures was tested once. For statistical analysis, we excluded 13 replicates with very low population density and/or infection prevalence (<10%), which prevented accurate estimation of dispersal of infected individuals. Dispersal was not significantly affected by assay date (χ^22^ = 2.56, p > 0.25), and this factor was therefore omitted from further analysis.

### Parasite life-history traits

#### Infectivity

On day 4 p.i., we estimated infection prevalence in the 30 inoculated cultures, using LAO fixation of ≈20 individuals, as described above. This measurement describes ‘parasite infectivity', i.e., the capacity to successfully establish infections (64).

#### Epidemiology and parasite development

From day 6 to 13 p.i., we tracked population density and infection prevalence in the 90 replicate cultures, using a blocked sliding window (day 6-8, 11-13) such that each parasite × host genotype combination was measured once per day. As infections developed, we also tracked changes in the proportion of infectious hosts, when reproductive forms are converted into infective spores. These data were used for the fitting of an epidemiological model (see below).

Furthermore, we specifically compared core and front parasites for their levels of infectiousness (= proportion of infectious hosts) between day 6 and day 11 p.i.. This time window describes the timing and level of investment into horizontal transmission by the initial cohort of infected hosts (72).

#### Virulence

Early after inoculation of the initial 30 assay cultures (day 4 p.i.), we isolated single infected and uninfected individuals from each culture and let them multiply for 9 days in 2-mL tubes under permissive common-garden conditions. From these small monoclonal cultures, we started the virulence assay by placing single individuals in PCR tubes filled with 200 μL of medium. We assessed cell division on day 2 and 3 (visual inspection), on day 10 (from 50-μL samples) and on day 20 from the total volume. A total of 645 replicates were set up, with 8-12 infected and uninfected replicates from each of 28 of the 30 assay cultures. For further details, see SI 2 and Fig. S3.

#### Swimming behaviour

From the above monoclonal lines, we placed 200-μL samples (containing 10-20 individuals) on a microscope slide and recorded individual movement trajectories using a Perfex SC38800 camera (15 frames per second; duration 10 s). For each sample, the mean net swimming speed and swimming tortuosity (standard deviation of the turning angle distribution, describing the extent of swimming trajectory change) was determined, using the BEMOVI package (75). This assay was performed for infected and uninfected monoclonal lines from 29 assay cultures, with 1-2 samples per monoclonal line (106 replicates in total).

### Statistical analysis

Statistical analyses were performed in R (ver. 3.3.3; R Development Core Team, available at www.r-project.org) and in JMP (SAS Institute Inc. (2018) JMP®, Version 14, N.C.). To analyse variation in parasite traits, we used generalised linear mixed-effect models (76). Binomial error structure and logit link were used for analysis of infected host dispersal (proportion dispersers), infectivity (proportion infected individuals on day 4 p.i.), and horizontal transmission investment (proportion infectious hosts day 6-11 p.i.). Normal error structure was used for analysis of swimming speed and tortuosity. For the virulence assay, we analysed variation in host division (= maximum cell density per replicate; Poisson error structure and log link) and survival (= replicate alive / dead on day 20; binomial error structure and logit link).

In all analyses, parasite selection treatment (front vs core) was taken as a fixed effect and host genotype and parasite selection line identity as random factors. Day p.i. was integrated as a covariate in the analysis of infectiousness. In the virulence analyses, replicate infection status (infected / uninfected) was included as a fixed factor. Analysis of variance (type II) was used to test for significance of fixed effects (car package; Fox and Weisberg, 2018). In complementary comparisons, we used ANOVA model predictions (and their variance) for core and front treatments to establish predictive distributions of the front-core difference (e.g. Fronhofer *et al.*, 2017). For these distributions, we calculated the mean difference and confidence intervals. Finally, we performed multiple regressions (path analysis) to assess how infected host dispersal was affected by the following traits: horizontal transmission investment (HTI, area under the curve of the proportion of infectious hosts from day 6 - 11 p.i.), virulence (infected host division) and swimming behaviour (tortuosity and net swimming speed). This analysis was based on trait means for 25 combinations of parasite selection line and host assay genotype. To meet assumptions of normality, certain trait means were transformed (log2 for host division, arcsine for dispersal, square root for HTI). To correct for overall effects of host genotype, we first fitted univariate analyses for each trait, and then performed the regressions on the residuals. Standardised beta regression coefficients were taken as path coefficients.

### Epidemiological model fitting

We fitted a simple epidemiological model to the above population density and infection prevalence data recorded in our assay replicate cultures (day 6-13 p.i.). The aim was to obtain additional independent estimates of parasite parameters (Table 1), namely virulence, but also the transmission parameter or latency time, i.e. the time to onset of production of infectious forms (Rosenbaum et al., 2019).

**Table 1.** Model parameters, their signification and the priors used for fitting.

| Parameters | Term | Priors |
| --- | --- | --- |
| $b$ | Birth rate | Posteriors from fitting eq. (1) on non-infected population data |
| $d$ | Death rate | |
| $\alpha$ | Intraspecific competition coefficient | |
| $\nu$ | Virulence (decrease in $b$) | Uniform (0, 1) |
| $\beta$ | Horizontal transmission rate | Lognormal (-5, 0.9) |
| $\gamma$ | Vertical transmission rate | Uniform (0.5, 1) |
| $\tau$ | Latency time | Uniform (144, 240) |

#### Model structure

We model the density of uninfected (S) and infected (I) hosts using ordinary differential equations (ODEs). In the absence of parasites, we consider that uninfected *Paramecium* growth follows the continuous time version of the Beverton-Holt model (49).

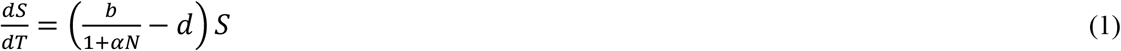

where *b* is the birth rate, *d* the death rate and *α* the competition term. *N* is the total number of individuals (*S +I*), which is equal to *S* in the absence of the parasite. In the presence of infected individuals, uninfected individuals become infected at a rate proportional to the number of infected and uninfected individuals at a rate *β*:

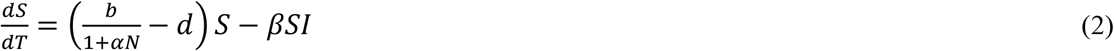

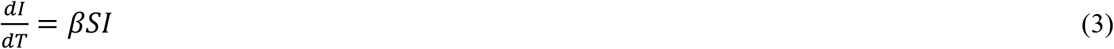

Moreover, infected individuals also display Beverton-Holt dynamics, but their birth rate can be decreased, hence we multiply *b* by a term *(1 – v)*, where *v* is the virulence of the parasite:

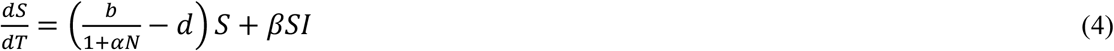

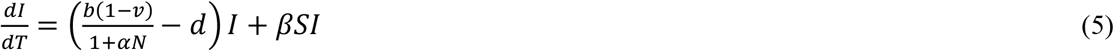

Finally, vertical transmission of the parasite is not necessarily 100%, and some of the *Paramecium* “born” from infected individuals could be free of parasites due to incomplete vertical transmission. We name *γ* the proportion of successful vertical transmission:

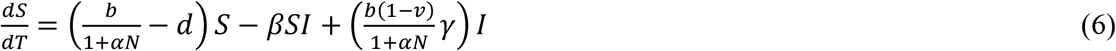

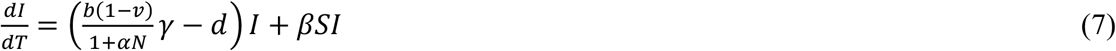

Since the majority of infected individuals were not yet producing infectious forms at the beginning of the time series, we added another parameter, *τ*, which is the latency before an infected individual becomes infectious (i.e., capable of horizontal transmission):

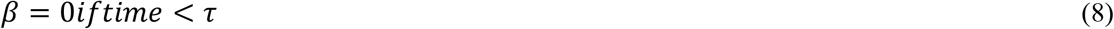

#### Model Fitting

We fitted the epidemiological model to the data using Bayesian inference and the *rstan* R package (version 2.19.2). Using data from previous experiments (O. Kaltz, unpublished data), we first fitted the Beverton-Holt model (Eq. 1) to growth curves of uninfected populations to estimate the distributions of *b, d* and *α* for each host genotype. These distributions were used as priors for fitting the full model (Eq. 6, 7, 8) on infection data. The model was fitted separately for each of the six combinations of host genotype and parasite selection treatment. For simplicity, we fitted a single set of parameters (*b, d, α, b, β, γ, τ*) over the different selection lines (with different initial conditions fitted over each line). Priors distributions can be found in Table 1. Apart from *b*, *d* and *α*, we used lowly informative priors that largely encompass expected values (*v* and *γ* priors are uniform over possible values, *τ* prior is uniform over previously observed latency values, *β* prior follows a lognormal distribution an order of magnitude wider than expected values). Fits were realized using the No U-Turn Sampler (NUTS) with default rstan values and multiple chains (three chains per fit, each of total length: 15 000 and warm-up length: 5 000).

## Acknowledgements

This work was supported by the 2019 Godfrey Hewitt mobility award granted to LN by ESEB, and by the Swiss National Science Foundation (grant no. P2NEP3_184489) to GZ. This is publication ISEM-YYYY-XXX of the Institut des Sciences de l'Evolution.

## Author contributions

OK, LN and GZ conceived the study. OK, LN, GZ, MH, and EAF helped design the experiments. LN, GZ, CGB and OK performed the experimental work., LN, GZ, MH, OK and EAF performed the statistical analysis. EAF, CS and OK developed, and CS analysed the epidemiological model. All authors interpreted the results and contributed to the writing of the manuscript.

## Data accessibility

If the manuscript is successfully accepted for publication, data will be available at Dryad or Figshare.

## Supplementary Information

### SI 1: Long-term selection protocol

Dispersal arenas consisted of two 14 mL plastic tubes (the core and front patch, Fig. S1) interconnected by a 5 cm silicon tube of 0.6 mm inner diameter (corridor). The protocol for a dispersal event is as follows. Step 1: We first fill the entire arena with 9.5 mL of fresh growth medium, then block the corridor with a clamp. Step 2: one of the two tubes (“core patch”) is topped up to 13 mL with 8 mL of medium containing cultures of *Paramecium caudatum (‘Paramecium’, hereafter)*, while the other tube (“front patch”) is topped up to 13 mL with medium only. Step 3: we remove the clamp for 3 h allowing the *Paramecium* to actively disperse from core to front patch or to stay in the core patch. Step 4: after blocking the corridor again, population densities in core and front patches are determined from 200-μL samples, with the number of individuals counted under a dissecting microscope. From these counts we can estimate the dispersal rate.

**Figure S1.**
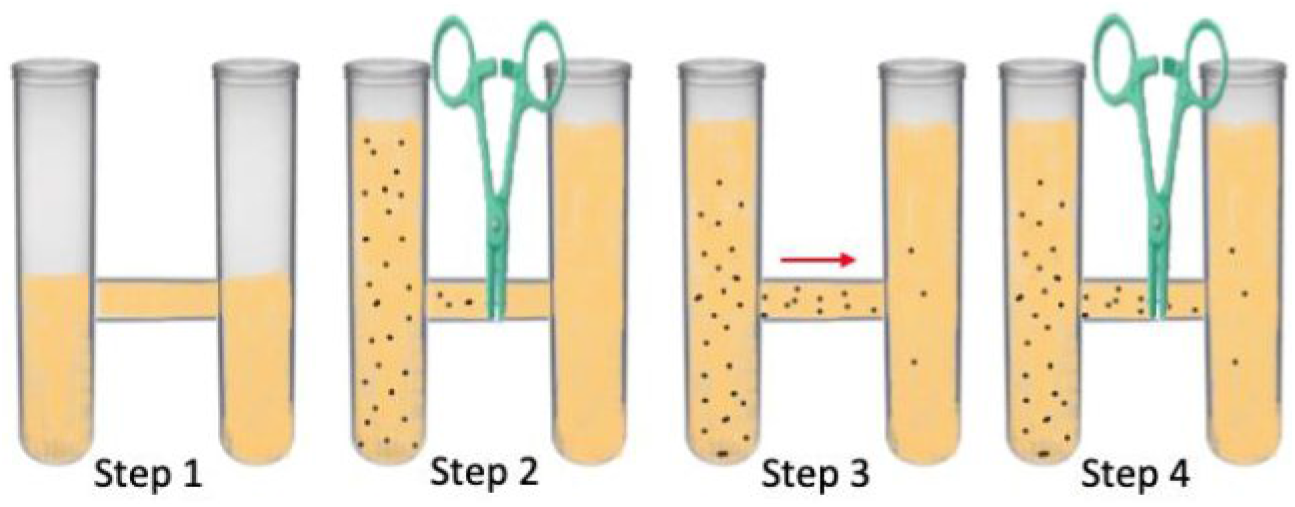
Protocol for setting up dispersal events in 2-patch dispersal arena. Step 1: filling dispersal arean up until dispersal corridor with fresh growth medium. Step 2: block corridor with a clamp and add cultures of *Paramecium* to patch 1 and fresh media to patch 2. Step 3: remove clamp and allow for acrive dispersal from patch 1 to patch 2 during a three-hour period. Step 4: block dispersal corridor and assess population densities and infection status in patch 1 and 2.

Using a mix of parasite inocula we infected naïve Paramecium of the 63D genotype to set up five front selection and five core selection lines (see main text for details on host and parasite origins). The long-term selection protocol is as follows (for illustration, see Fig. 1 in main text).

(i) In the front selection treatment, densities in the core patch prior to dispersal were set to c. 2000 individuals. After the 3 h of free dispersal, only *Paramecium* that dispersed to the front patch were maintained, transferred to a 50-mL plastic tube and cultured for one week in 20 mL of bacterised lettuce medium, before the next dispersal and selection event occurred. On average, c. 350 individuals dispersed at a given dispersal event. When we observed fewer than 100 dispersers, we topped up numbers to 100 by adding non-dispersers. Over the one week of culture, populations grew back to carrying capacity (c. 4-5 × 10^3^ individuals) and epidemiological processes were acting freely. Then a new episode of dispersal occurred, as described above.

(ii) In the core selection treatment, we followed the same protocol as above, but only *Paramecium* that stayed in the core patch were maintained and regrown in 20 mL of medium. Furthermore, numbers of transferred core individuals were adjusted so as to match those in the front selection treatment at the beginning of the one-week growth period.

After 30 dispersal/growth cycles, we extracted infectious forms from each selection line and used them to inoculate a new batch of 63D hosts. This was done to minimise effects of host evolution in the experiment and to continue parasite evolution on naïve hosts for another round of 25 cycles. Because carrying capacity in this new round (c. 3-4 × 10^3^ individuals) was lower than in the first 30 cycles, we relaxed the dispersal protocol. Instead of adjusting the density prior to dispersal, we topped up the core patch with the maximum possible volume (8 mL; step 2, see above) to ensure that at least 10^3^ were placed in the tube.

### SI 2: Assay details

After inoculation of three naive host genotypes (63D, C023, C173) with evolved core and front parasites, a series of assays were performed over a 3-week period. In this section, we present an overview of the timing of the assays (Table S1) and further details on the experimental protocols.

**Table S1.** Timing of measurements for the different parasite traits in the assay. The assay was started by inoculating naive cultures of three host genotypes with parasites from 5 core and 5 front selection treatments (30 combinations in total). Timing of each assay is expressed in days post inoculation (p.i.). On day 4 p.i., the initial 30 inoculated replicates were split into three technical assay replicates, giving a total of 90 assay replicates. For certain traits, not all replicate cultures were available, or data are missing. The ‘Number’ columns indicate the number of combinations and replicate assay cultures available for each analysis. ‘Total n’ refers to the total number of data points in the analysis (raw data). *Paramecium* for the singleton assay were isolated from the initial 30 inoculated assay cultures. For details on trait measurements, see SI below and main text.

| Trait | Assay day (p.i.) | Number of host x parasite combinations (max. 30) | Number of assay replicate cultures (out of total) | Total n (raw data) | Comments |
| --- | --- | --- | --- | --- | --- |
| Infectivity | 4 | 30 | 30/30 | 30 |  |
| Infectiousness | 4 - 11 | 30 | 90/90 | 147 | First infectious hosts detected on day 6 p.i. |
| Infected dispersal | 13 - 19 | 27 | 75/90 | 75 | Exclude replicate cultures if low density / infection prevalence (<10%) |
| Virulence<br>(Host division and survival in singleton assay) | 13 - 23 | 28 | 28/30 | 605 | Infected and uninfected individuals isolated on day 4 p.i. and grown for 9 days under common-garden conditions prior to singleton assay |
| Movement<br>(tortuosity and swimming speed) | 14 | 29 | 29/30 | 106 | Video analysis performed on infected and uninfected populations isolated on day 4 p.i. |

#### Infectivity

##### Inoculum preparation

To extract infectious forms of *Holospora undulata* from an infected culture, the infected *Paramecium* are transferred to 50-mL Falcon tubes and centrifuged at 1500 g for 20 minutes. The supernatant is removed, and the concentrated individuals placed in 1.5 mL Eppendorf tubes containing 1 mm glass beads. Using a Qiagen TissueLyser they are then vortexed and crushed (1:45 minutes at 30 oscillation frequency) to liberate infectious forms of the parasite. The concentration of infectious forms is then determined at 200x magnification under a microscope (Leica DM LB2), using a hemocytometer.

##### Infectivity assay

Inocula of the evolved parasites selected for high or low dispersal (see Fig. S2; front and core, respectively) were prepared, as described above. Their capacity to establish infections (= infectivity) was tested on samples of unselected naïve hosts, represented by three genotypes: 63D, C023 and C173. Prior to inoculation, cultures of the three genotypes were concentrated by using mild centrifugation (15 min at 300 g). We placed ≈ 5 × 10^3^ cells of a given host genotype in 1.4 mL of bacterised medium in a 15 mL tube, to which we added the freshly prepared inoculum of a given evolved parasite line. Inoculum dose ranged from c. 0.3 - 1.4 × 10^3^ infective spores per μL (median: 1 × 10^3^), depending on the identity of parasite selection line. These doses represent an *ad libitum* administration; typically, infection success reaches a plateau for doses > 0.2 spores per μL (Fels et al 2008). In the present experiment, a preliminary logistic regression with binomial error structure (logit link) revealed no significant effect of spore dose on infection success (χ_1_^2^ = 10.812, p = 0.147).

### Dispersal assay

In the assay, we used 3-patch arenas (50-mL Falcon tubes; Fig. S3) instead of the usual 2-patch arena. These arenas allowed us to use larger numbers of *Paramecium* and, by letting them disperse from the middle tube into the two outer tubes, we increased the total number of dispersers. To further increase the resolution of dispersal estimates, we concentrated the cultures 12 h prior to the assay, by gentle centrifugation at 300 g for 15 minutes. For the dispersal assay, we first filled the 3-patch system with 20 mL of fresh medium, and then blocked the two corridors with clamps (see steps 1 and 2 in Fig. S1). We then added ≈2800 individuals to the middle tube and topped it up to 25 mL with fresh medium. The outer tubes were topped up to 25 mL with fresh medium only, and the corridors were then opened for 3 h to allow for free dispersal. At the end of the 3-h dispersal period, we determined population density and infection prevalence, for samples from the central tube (500 μl) and from the combined two outer tubes (3 mL). From these estimates we calculated the dispersal rate of infected hosts, as explained above. Of the 90 inoculated replicate assay cultures, 88 were tested (two tubes were found to be uninfected). For statistical analysis, we excluded 13 replicates with very low population density and/or low infection prevalence (<10%), which prevented accurate estimation of dispersal of infected individuals. We also assayed a small number of uninfected control cultures (three tubes per host genotype) for comparison with dispersal rates in infected tubes. For unknown reasons, these replicates showed considerable variation; we therefore regrouped these observations with measurements from previous experiments to inform on the general range of dispersal rates observed for these genotypes (see Fig. S4). We did not carry out formal statistical comparisons with infected dispersal rates observed in our assay.

**Figure S2.**
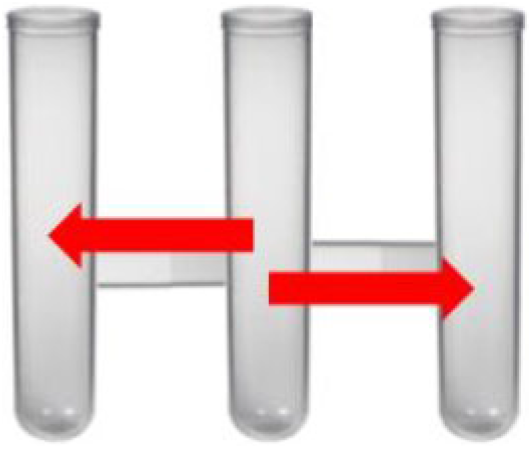
Linear 3-patch arenas used for the dispersal assays. *Paramecium* was added in the central tube and after the opening of the connections it was allowed to disperse (3h) in the outer tubes, as indicated by the red arrows. For filling protocol, see Fig. S1.

### Virulence assay

For the virulence assay, we isolated single infected and uninfected individuals from the replicate assay cultures on day 4 p.i and allowed them to multiply in 2-mL Eppendorf tubes with bacterised medium, under permissive common-garden conditions. After one week, we determined infection status of these monoclonal cultures (LAO fixation). On day 13 p.i., we started the assay by placing single individuals in PCR tubes filled with 200 μL of bacterised medium (Fig. S3). We checked tubes daily for presence or absence of live cells for 20 days. In addition, cell density was determined on day 2 and 3 (by counting cells through the plastic tubes), on day 10 (from 50-μL samples) and on day 20 (total volume). Except for 50 μL of medium added on day 10, no resources were supplied.

**Figure S3.**
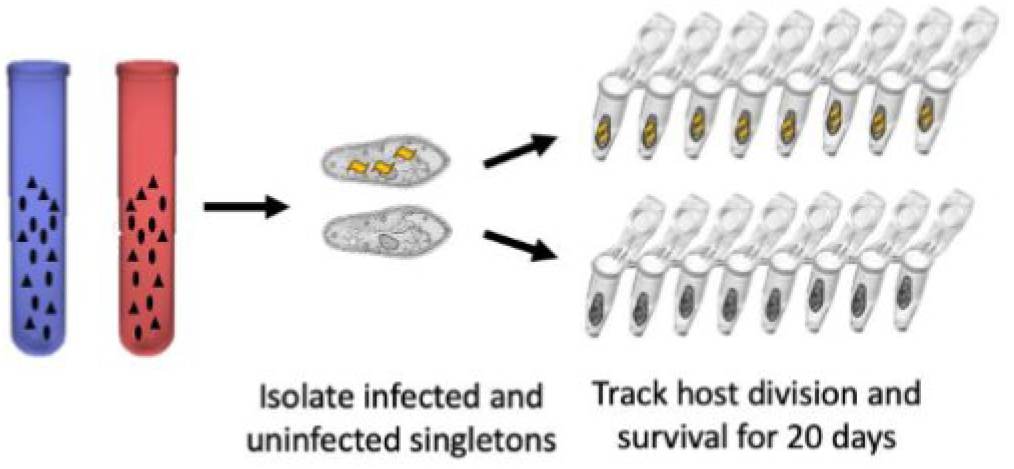
Protocol for virulence assay. On day 4 post inoculation of replicate cultures with front (ref) or core (blue) parasites, infected and uninfected individuals were isolated and allowed to grow as small monoclonal cultures. Single individuals were then placed PCR tubes containing 200 μL bacterised medium. Each combination of parasite selection line (n = 10) and host genotype (n = 3) was replicated 8-12 times.

The assay was performed for 28 of the 30 combinations of parasite selection line and host genotypes, with 8-12 infected singletons tested per combination. Of the 322 infected replicates, 17 died within less than 24 h (possibly due to transfer handling) and were excluded from statistical analysis. In the same way, excluded 23 early deaths of the 323 uninfected singletons.

### SI 3: Results

The assays of the evolved parasites were performed on naïve *Paramecium* representing three host genotypes (63D, C172, C023). The parasites had evolved on the 63D genotype in the selection experiment, but had never been exposed to any of the other two genotypes. In our main statistical analyses, we considered host genotype as a random effect, as the experimental tests were not specifically meant to address differences among the host genotypes. These results are summarised in Table S2.

**Table S2.**
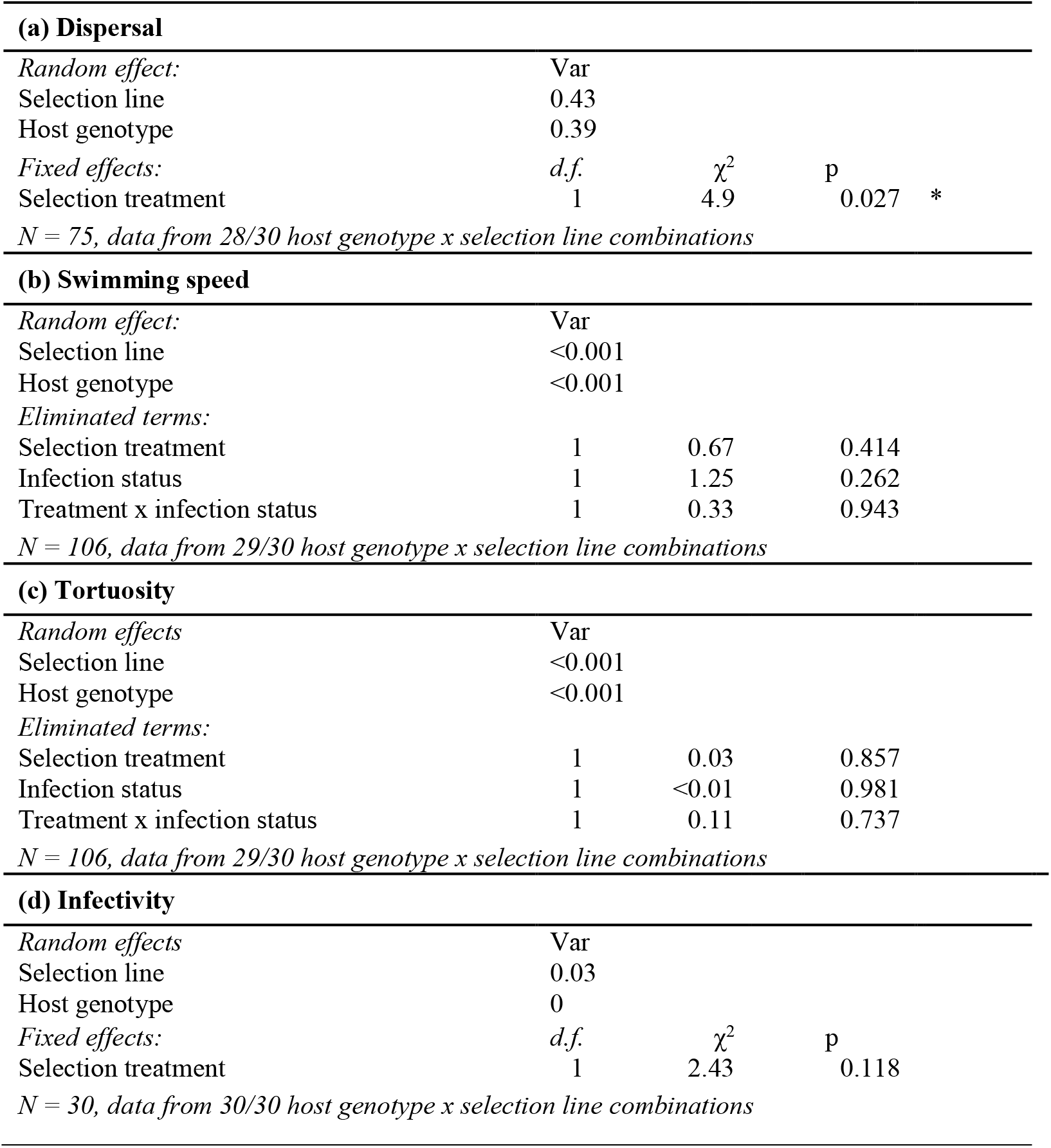
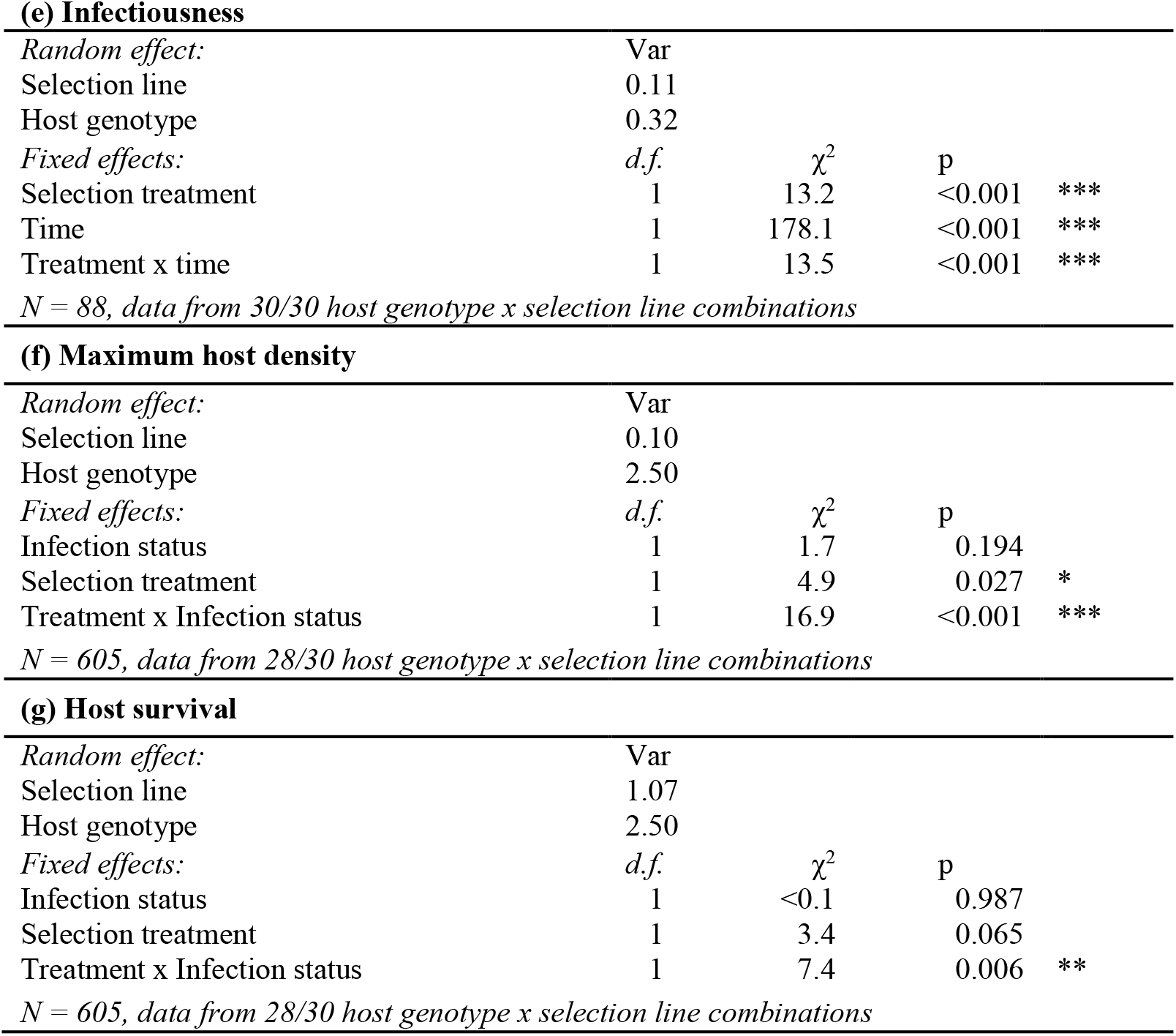
ANOVA results from GLMM models of all measured traits (dispersal, swimming speed and tortuosity, infectivity, infectiousness and virulence). Parasite selection line and host genotype were added as random factors, all other factors were considered fixed. We performed backward model simplification, with sequential removal on non-significant terms in the model. Terms in the “Fixed effects” category represent the minimal adequate model; sequentially removed terms appear in the “Eliminated terms” category. For each analysis, we indicate the number of replicates used in the analysis and the number of host genotype x parasite selection lines they represent (3 × 10 = 30 combinations inoculated, some of which were not available for certain analyses).

|  |  |  |  |  |
| --- | --- | --- | --- | --- |
| **(a) Dispersal** |  |  |  |  |
| *Random effect:* | Var |  |  |  |
| Selection line | 0.43 |  |  |  |
| Host genotype | 0.39 |  |  |  |
| *Fixed effects:* | *d.f.* | $\chi^2$ | p | |
| Selection treatment | 1 | 4.9 | 0.027 | * |
| *N = 75, data from 28/30 host genotype x selection line combinations* |  |  |  |  |
| **(b) Swimming speed** |  |  |  |  |
| *Random effect:* | Var |  |  |  |
| Selection line | <0.001 |  |  |  |
| Host genotype | <0.001 |  |  |  |
| *Eliminated terms:* |  |  |  |  |
| Selection treatment | 1 | 0.67 | 0.414 |  |
| Infection status | 1 | 1.25 | 0.262 |  |
| Treatment x infection status | 1 | 0.33 | 0.943 |  |
| *N = 106, data from 29/30 host genotype x selection line combinations* |  |  |  |  |
| **(c) Tortuosity** |  |  |  |  |
| *Random effects* | Var |  |  |  |
| Selection line | <0.001 |  |  |  |
| Host genotype | <0.001 |  |  |  |
| *Eliminated terms:* |  |  |  |  |
| Selection treatment | 1 | 0.03 | 0.857 |  |
| Infection status | 1 | <0.01 | 0.981 |  |
| Treatment x infection status | 1 | 0.11 | 0.737 |  |
| *N = 106, data from 29/30 host genotype x selection line combinations* |  |  |  |  |
| **(d) Infectivity** |  |  |  |  |
| *Random effects* | Var |  |  |  |
| Selection line | 0.03 |  |  |  |
| Host genotype | 0 |  |  |  |
| *Fixed effects:* | *d.f.* | $\chi^2$ | p | |
| Selection treatment | 1 | 2.43 | 0.118 |  |
| *N = 30, data from 30/30 host genotype x selection line combinations* |  |  |  |  |

Table S2 (cont.) **(e) Infectiousness**
|  |  |  |  |  |
| --- | --- | --- | --- | --- |
| *Random effect:* | Var |  |  |  |
| Selection line | 0.11 |  |  |  |
| Host genotype | 0.32 |  |  |  |
| *Fixed effects:* | *d.f.* | $\chi^2$ | p | |
| Selection treatment | 1 | 13.2 | <0.001 | *** |
| Time | 1 | 178.1 | <0.001 | *** |
| Treatment x time | 1 | 13.5 | <0.001 | *** |
| **(f) Maximum host density** |  |  |  |  |
| *Random effect:* | Var |  |  |  |
| Selection line | 0.10 |  |  |  |
| Host genotype | 2.50 |  |  |  |
| *Fixed effects:* | *d.f.* | $\chi^2$ | p | |
| Infection status | 1 | 1.7 | 0.194 |  |
| Selection treatment | 1 | 4.9 | 0.027 | * |
| Treatment x Infection status | 1 | 16.9 | <0.001 | *** |
| **(g) Host survival** |  |  |  |  |
| *Random effect:* | Var |  |  |  |
| Selection line | 1.07 |  |  |  |
| Host genotype | 2.50 |  |  |  |
| *Fixed effects:* | *d.f.* | $\chi^2$ | p | |
| Infection status | 1 | <0.1 | 0.987 |  |
| Selection treatment | 1 | 3.4 | 0.065 |  |
| Treatment x Infection status | 1 | 7.4 | 0.006 | ** |

*N* = 605, data from 28/30 host genotype x selection line combinations

To inspect the generality of selection treatment effects, we also performed analyses, treating host genotype as a fixed effect and including the selection treatment × host genotype interaction (Table S3). They are accompanied by a more detailed presentation of trait expression, with separate panels for each host genotype (Fig. S4–S10). These additional analyses showed that, with few exceptions, differences between core and front parasites were consistent across the different hosts (i.e., treatment × host interactions were non-significant), even though in some cases more or less pronounced. For details of trait definitions and measurements, see main text.

**Table S3.**
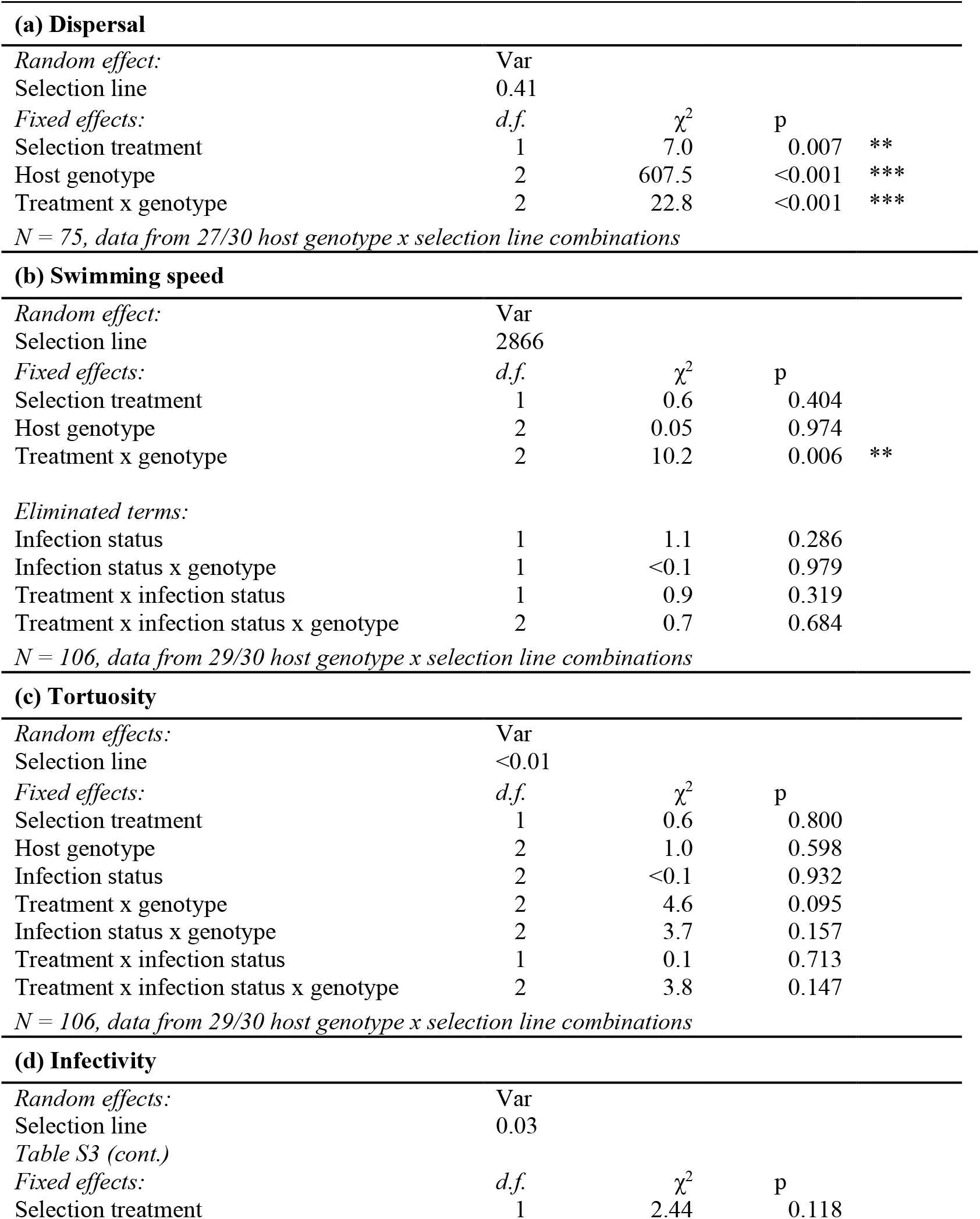
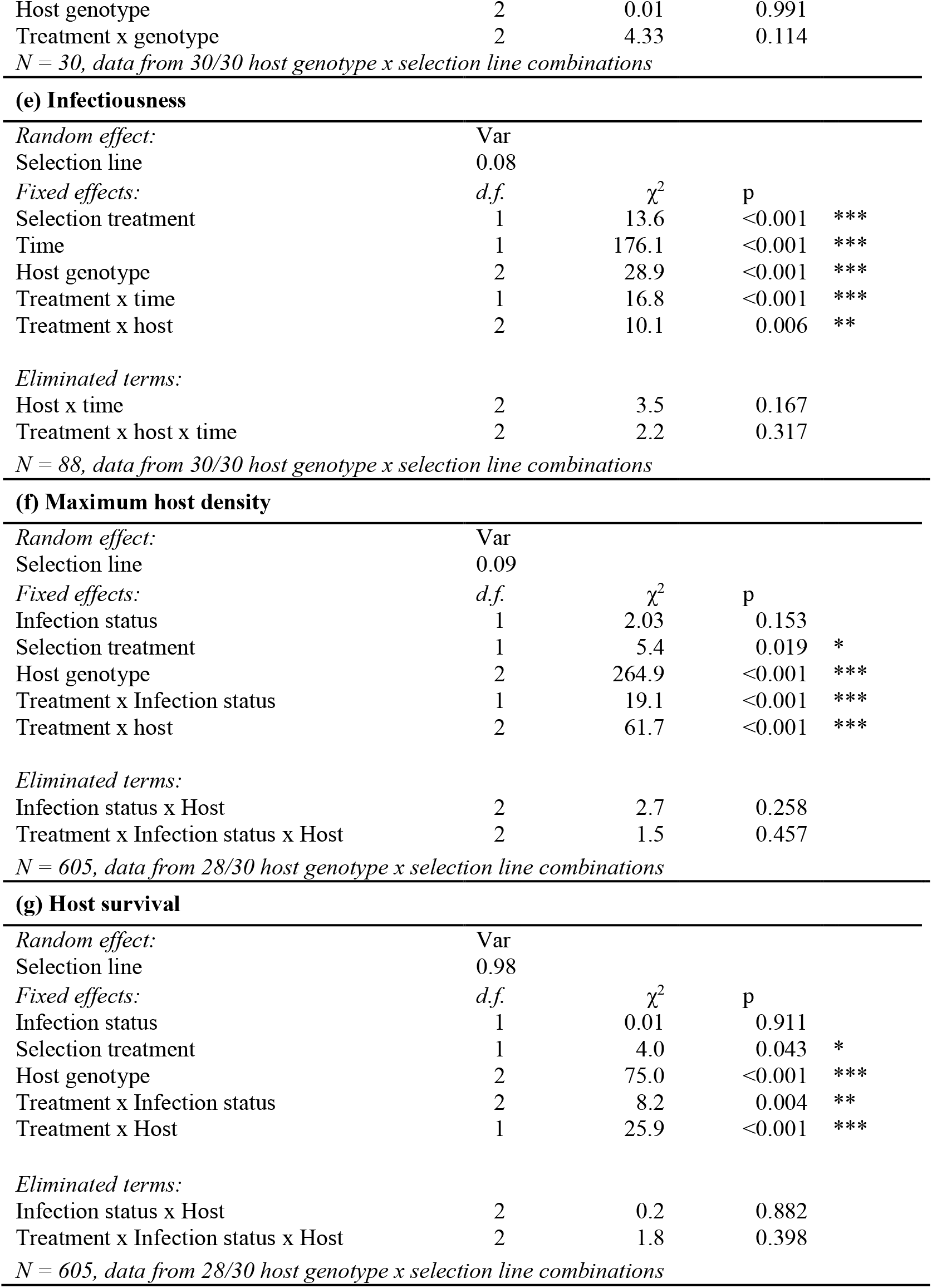
ANOVA results from GLMM models of all measured traits (dispersal, swimming speed and tortuosity, infectivity, infectiousness and virulence). Parasite selection line was treated as a random effect, all other factors were fixed effects. We performed backward model simplification, with non-significant terms sequentially removed from the model (“eliminated terms”). For each analysis, we indicate the number of replicates and the number of host genotype x parasite selection lines they represent (3 × 10 = 30 combinations inoculated, some of which were not available for certain analyses).

|  |  |  |  |  |  |
| --- | --- | --- | --- | --- | --- |
| **(a) Dispersal** |  |  |  |  |  |
| *Random effect:* | Var |  |  |  |  |
| Selection line | 0.41 |  |  |  |  |
| *Fixed effects:* | *d.f.* | $\chi^2$ | p | | |
| Selection treatment | 1 | 7.0 | 0.007 | ** |  |
| Host genotype | 2 | 607.5 | <0.001 | *** |  |
| Treatment x genotype | 2 | 22.8 | <0.001 | *** |  |
| *N = 75, data from 27/30 host genotype x selection line combinations* |  |  |  |  |  |
| **(b) Swimming speed** |  |  |  |  |  |
| *Random effect:* | Var |  |  |  |  |
| Selection line | 2866 |  |  |  |  |
| *Fixed effects:* | *d.f.* | $\chi^2$ | p | | |
| Selection treatment | 1 | 0.6 | 0.404 |  |  |
| Host genotype | 2 | 0.05 | 0.974 |  |  |
| Treatment x genotype | 2 | 10.2 | 0.006 | ** |  |
| *Eliminated terms:* |  |  |  |  |  |
| Infection status | 1 | 1.1 | 0.286 |  |  |
| Infection status x genotype | 1 | <0.1 | 0.979 |  |  |
| Treatment x infection status | 1 | 0.9 | 0.319 |  |  |
| Treatment x infection status x genotype | 2 | 0.7 | 0.684 |  |  |
| *N = 106, data from 29/30 host genotype x selection line combinations* |  |  |  |  |  |
| **(c) Tortuosity** |  |  |  |  |  |
| *Random effects:* | Var |  |  |  |  |
| Selection line | <0.01 |  |  |  |  |
| *Fixed effects:* | *d.f.* | $\chi^2$ | p | | |
| Selection treatment | 1 | 0.6 | 0.800 |  |  |
| Host genotype | 2 | 1.0 | 0.598 |  |  |
| Infection status | 2 | <0.1 | 0.932 |  |  |
| Treatment x genotype | 2 | 4.6 | 0.095 |  |  |
| Infection status x genotype | 2 | 3.7 | 0.157 |  |  |
| Treatment x infection status | 1 | 0.1 | 0.713 |  |  |
| Treatment x infection status x genotype | 2 | 3.8 | 0.147 |  |  |
| *N = 106, data from 29/30 host genotype x selection line combinations* |  |  |  |  |  |
| **(d) Infectivity** |  |  |  |  |  |
| *Random effects:* | Var |  |  |  |  |
| Selection line | 0.03 |  |  |  |  |
| *Table S3 (cont.)* |  |  |  |  |  |
| *Fixed effects:* | *d.f.* | $\chi^2$ | p | | |
| Selection treatment | 1 | 2.44 | 0.118 |  |  |

|  |  |  |  |
| --- | --- | --- | --- |
| Host genotype | 2 | 0.01 | 0.991 |
| Treatment x genotype | 2 | 4.33 | 0.114 |
| *N = 30, data from 30/30 host genotype x selection line combinations* |  |  |  |

(e) Infectiousness
|  |  |  |  |  |
| --- | --- | --- | --- | --- |
| *Random effect:* | Var |  |  |  |
| Selection line | 0.08 |  |  |  |
| *Fixed effects:* | *d.f.* | $\chi^2$ | p | |
| Selection treatment | 1 | 13.6 | <0.001 | *** |
| Time | 1 | 176.1 | <0.001 | *** |
| Host genotype | 2 | 28.9 | <0.001 | *** |
| Treatment x time | 1 | 16.8 | <0.001 | *** |
| Treatment x host | 2 | 10.1 | 0.006 | ** |
| *Eliminated terms:* |  |  |  |  |
| Host x time | 2 | 3.5 | 0.167 |  |
| Treatment x host x time | 2 | 2.2 | 0.317 |  |

(f) Maximum host density
|  |  |  |  |  |
| --- | --- | --- | --- | --- |
| *Random effect:* | Var |  |  |  |
| Selection line | 0.09 |  |  |  |
| *Fixed effects:* | *d.f.* | $\chi^2$ | p | |
| Infection status | 1 | 2.03 | 0.153 |  |
| Selection treatment | 1 | 5.4 | 0.019 | * |
| Host genotype | 2 | 264.9 | <0.001 | *** |
| Treatment x Infection status | 1 | 19.1 | <0.001 | *** |
| Treatment x host | 2 | 61.7 | <0.001 | *** |
| *Eliminated terms:* |  |  |  |  |
| Infection status x Host | 2 | 2.7 | 0.258 |  |
| Treatment x Infection status x Host | 2 | 1.5 | 0.457 |  |

(g) Host survival
|  |  |  |  |  |
| --- | --- | --- | --- | --- |
| *Random effect:* | Var |  |  |  |
| Selection line | 0.98 |  |  |  |
| *Fixed effects:* | *d.f.* | $\chi^2$ | p | |
| Infection status | 1 | 0.01 | 0.911 |  |
| Selection treatment | 1 | 4.0 | 0.043 | * |
| Host genotype | 2 | 75.0 | <0.001 | *** |
| Treatment x Infection status | 2 | 8.2 | 0.004 | ** |
| Treatment x Host | 1 | 25.9 | <0.001 | *** |
| *Eliminated terms:* |  |  |  |  |
| Infection status x Host | 2 | 0.2 | 0.882 |  |
| Treatment x Infection status x Host | 2 | 1.8 | 0.398 |  |

*N = 605, data from 28/30 host genotype x selection line combinations*

#### Dispersal

**Figure S4.**
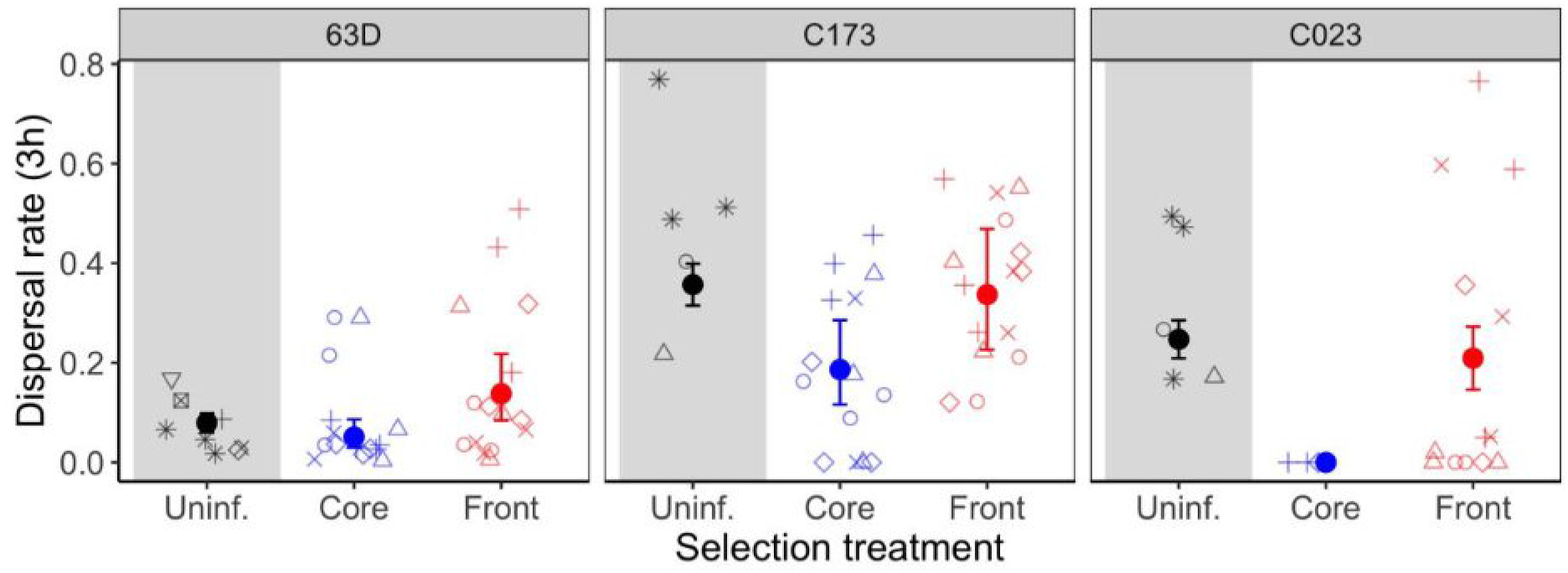
Dispersal rate (3 hr) of three naïve *Paramecium* genotypes (63D, C173, C023), infected with core parasites (blue) or front parasites (red). Data of uninfected replicates (black) from this assay and other experiments indicate the typical range of dispersal for each genotype (low for 63D, higher for C173 and C023). Filled circles and error bars represent means ± 95% confidence interval. Open symbols represent raw data. Different symbols refer to different parasite selection lines (5 lines per selection treatment).

#### Net swimming speed

**Figure S5.**
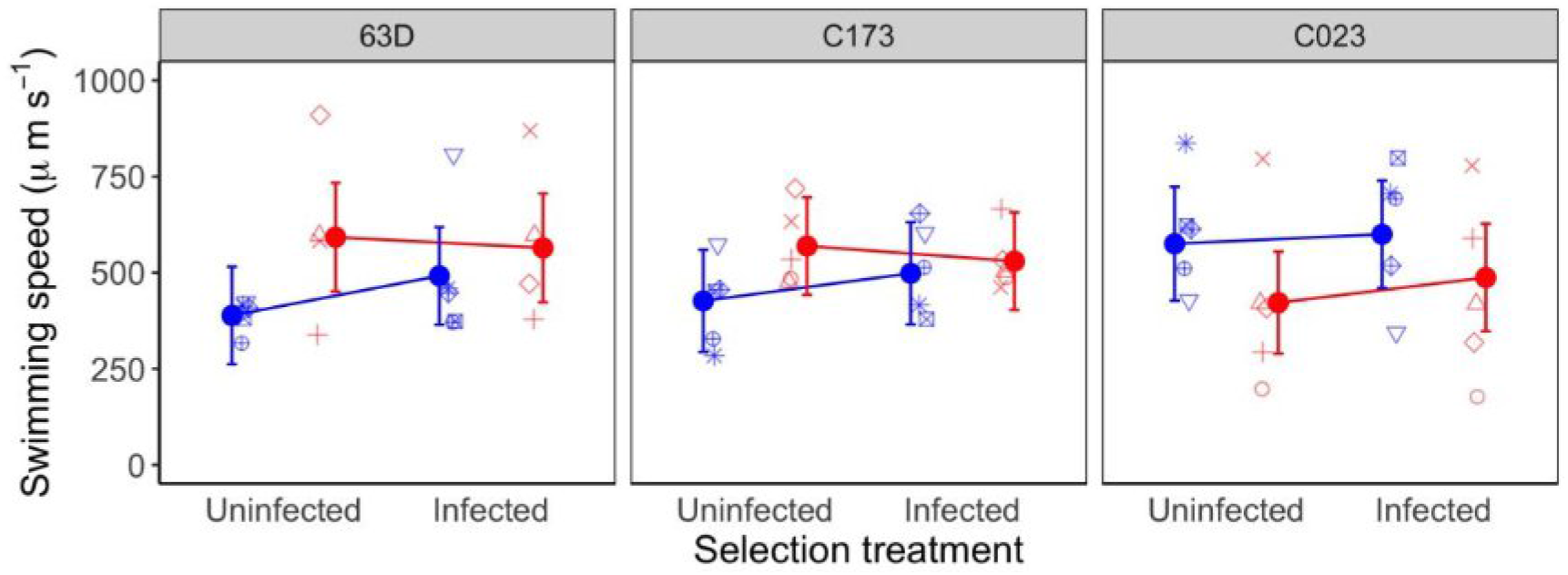
Net swimming speed (μm s^−1^) of three naïve *Paramecium* genotypes (63D, C173, C023), when uninfected or infected with core parasites (blue) or front parasites (red). Filled circles and error bars represent mean model predictions ± 95% confidence interval. Open symbols represent means over two technical replicates. Different symbols refer to different parasite selection lines (5 lines per selection treatment).

#### Tortuosity - trajectory variation

**Figure S6.**
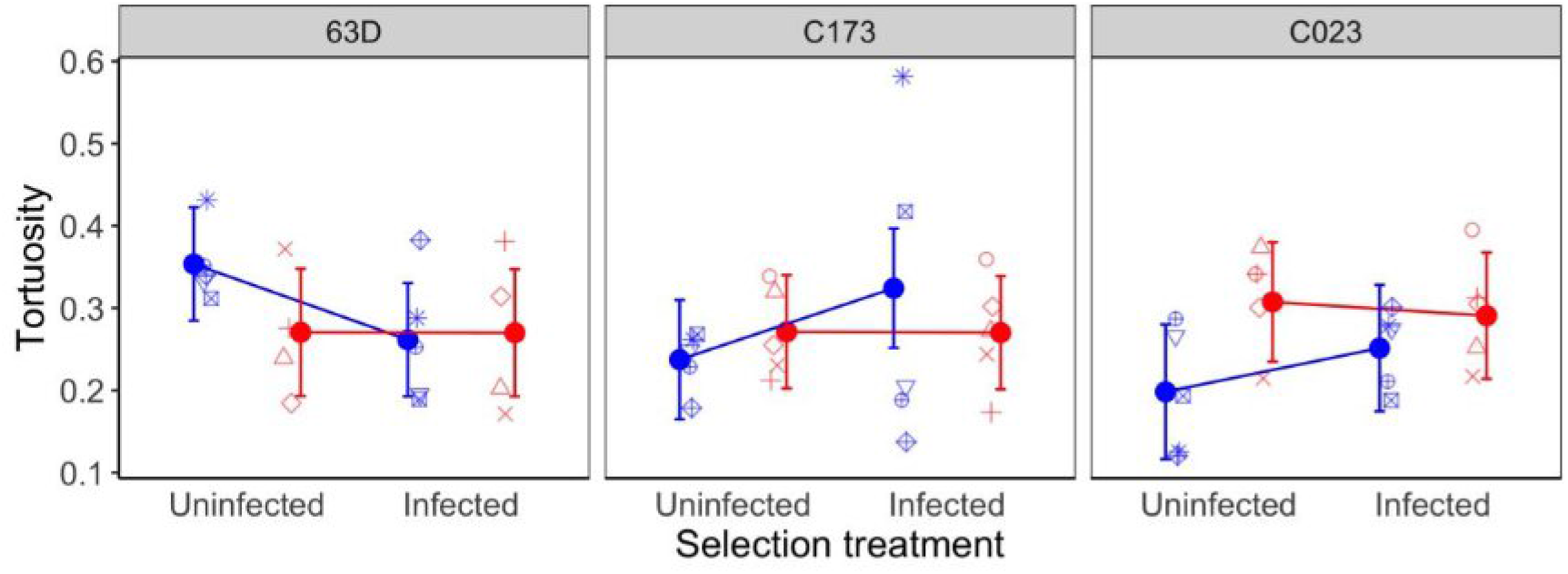
Swimming trajectory changes (tortuosity = standard deviation of turning angle distribution) of three naïve *Paramecium* genotypes (63D, C173, C023), when uninfected or infected with core parasites (blue) or front parasites (red). Filled circles and error bars represent mean model predictions ± 95% confidence interval. Open symbols represent means over two technical replicates. Different symbols refer to different parasite selection lines (5 lines per selection treatment).

#### Infectivity

**Figure S7.**
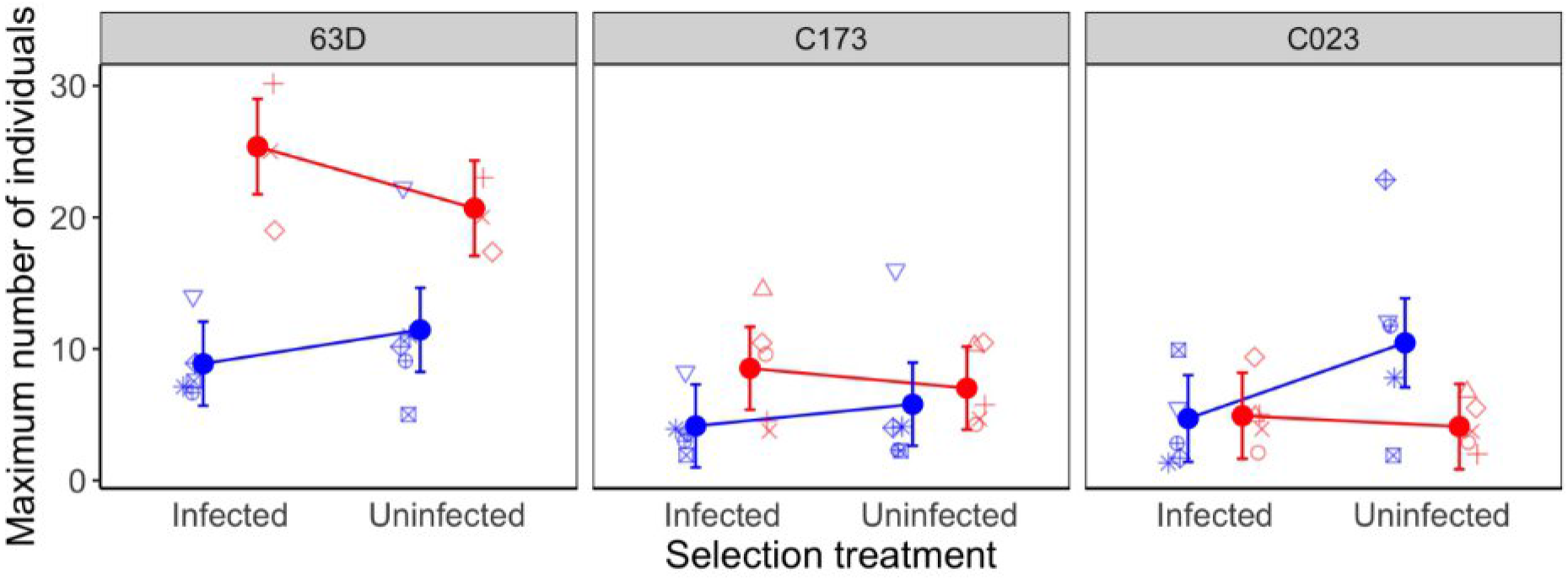
Proportion of infected individuals in assay cultures of three naïve *Paramecium* genotypes (63D, C173, C023), measured on day 4 post inoculation with core parasites (blue) or front parasites (red). Filled circles and error bars represent mean model predictions ± 95% confidence interval. Open symbols represent raw data. Different symbols refer to different parasite selection lines (5 lines per selection treatment). Note that these data were obtained before to splitting each culture into the 3 technical replicates (see Material and Methods in main text).

#### Infectiousness

**Figure S8.**
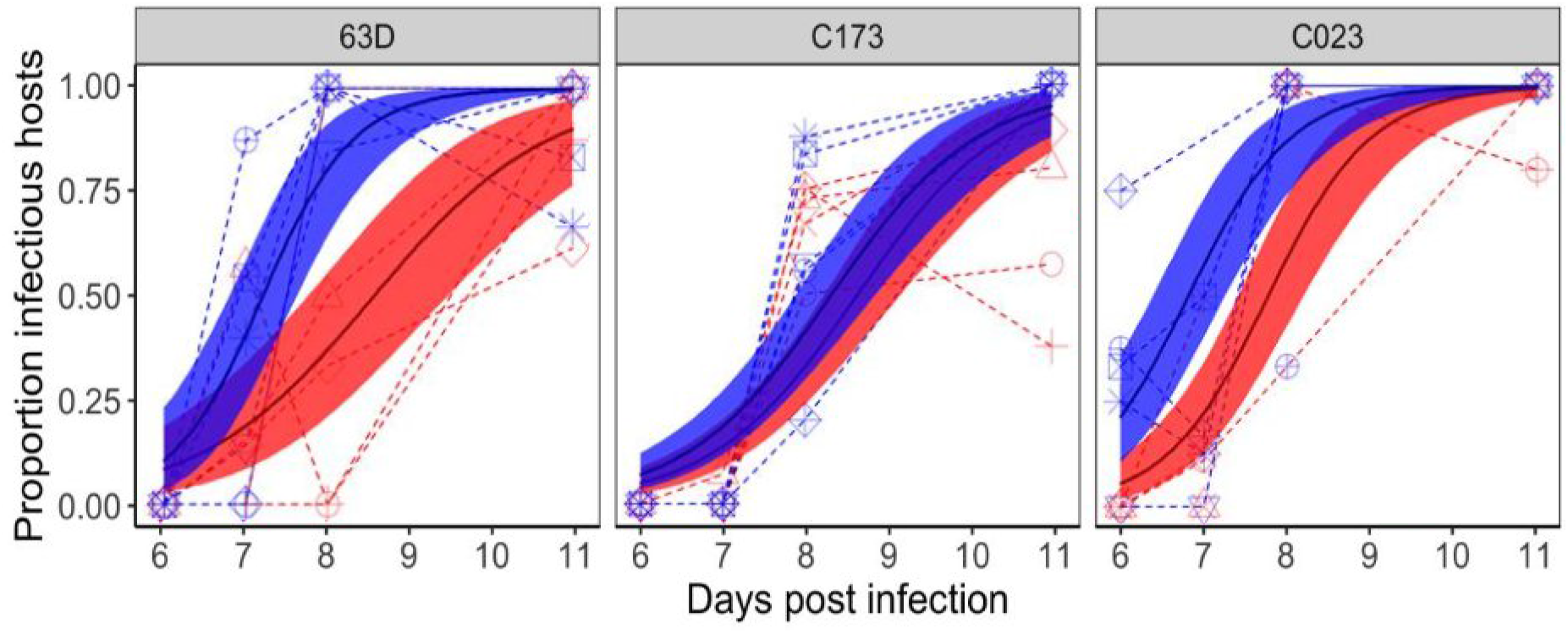
Infectiousness (proportion of infectious hosts, i.e., individuals host carrying infective spores) on day 6, 7, 8 and 11 post inoculation (p.i.), shown for three naïve *Paramecium* genotypes (63D, C173, C023) infected with core parasites (blue) or front parasites (red). Filled circles and error bars represent mean model predictions ± 95% confidence interval. Solid lines represent mean trajectories, stippled lines represent raw data trajectories for different parasite selection lines (note that certain trajectories are superimposed). Day 6 p.i. was the first day of detection of infectious hosts.

#### Virulence (I): Host division

**Figure S9.**
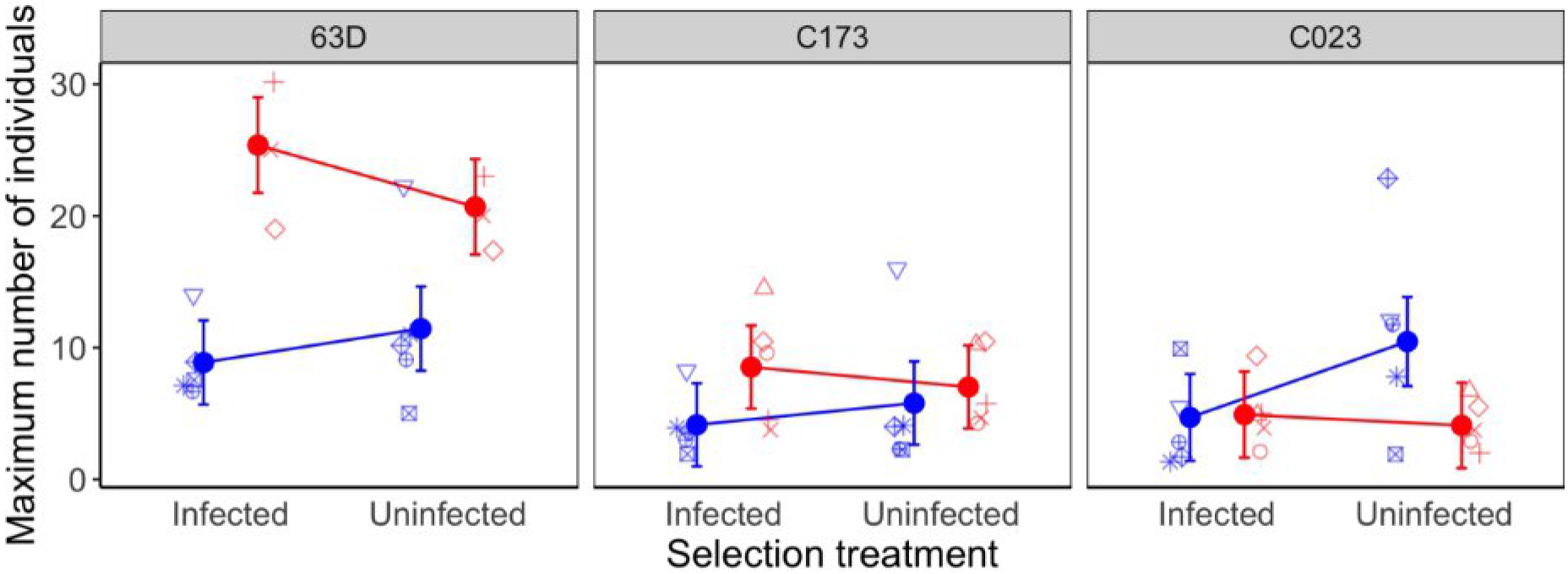
Maximum cell density of three naïve *Paramecium* genotypes (63D, C173, C023), when uninfected or infected with core parasites (blue) or front parasites (red). Data obtained in a 20-day assay, starting from single individuals, where maximum densities were generally reached on day 10, although some samples died sooner and thus peaked earlier. Filled circles and error bars represent mean model predictions ± 95% confidence interval. Open symbols represent means over c. 10 technical replicates, on average. Different symbols refer to different parasite selection lines (5 lines per selection treatment).

#### Virulence (II): Host survival

**Figure S10.**
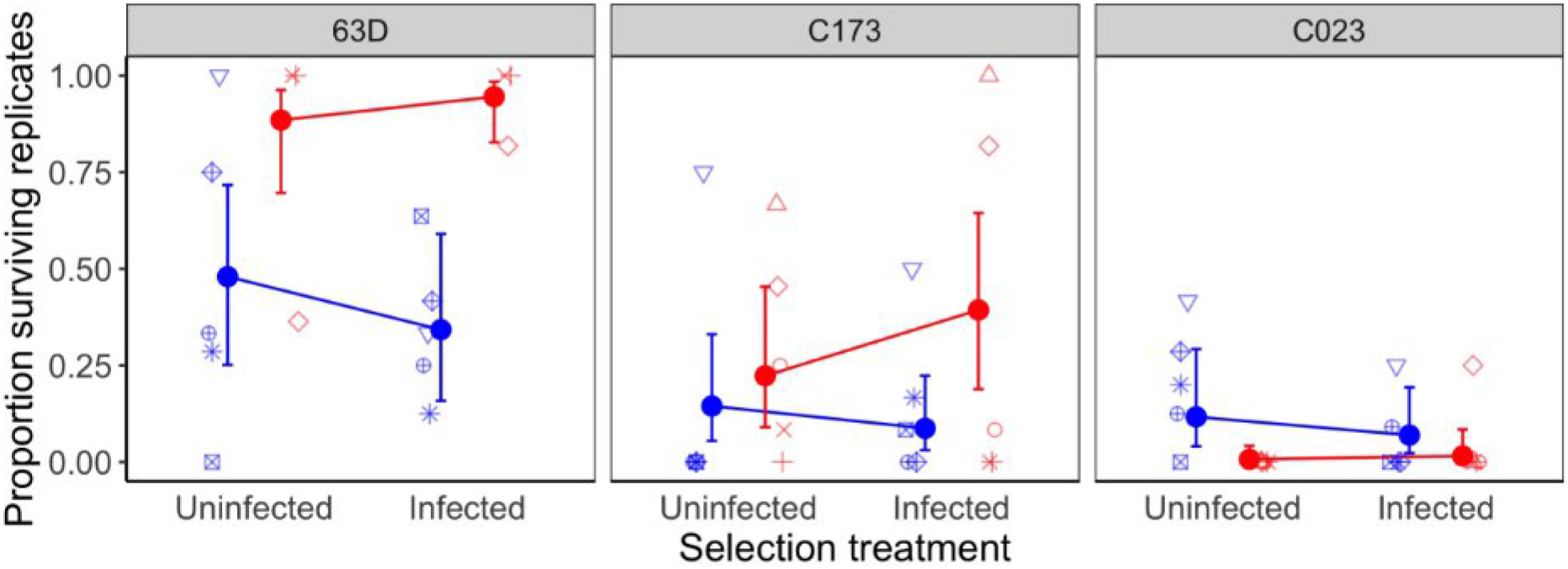
Mortality of three naïve *Paramecium* genotypes (63D, C173, C023), when uninfected or infected with core parasites (blue) or front parasites (red). Data show the proportion of replicates found extinct on day 20 in an assay starting from single individuals. Filled circles and error bars represent mean model predictions ± 95% confidence interval. Open symbols represent the observed proportions for the different parasite selection lines (5 lines per selection treatment).

### SI 4: Epidemiological model fits

**Figure S11.**
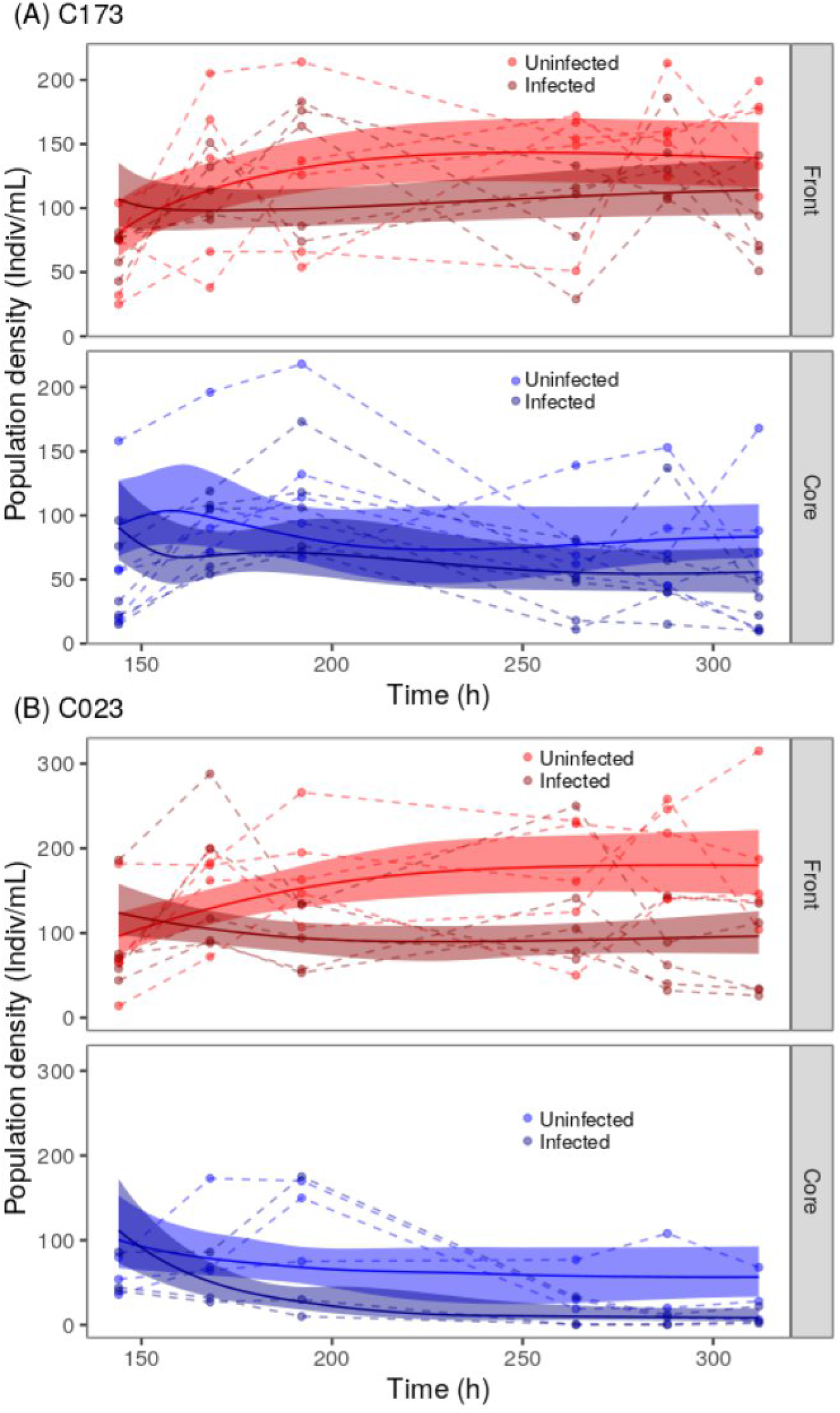
Fit of the epidemiological model (equations 6-8) to infected and uninfected host density time series data in the core and front treatment for host genotype (A) C173 and (B) C023. Dashed lines show observed density trajectories, solid lines and shaded areas represent posterior model predictions.

## Notes

### Competing Interest Statement

The authors have declared no competing interest.

### Summary of Updates

Analysis and discussion clarified, Figures updated; Supplementary section updated

